# Load adaptation of endocytic actin networks

**DOI:** 10.1101/2020.04.05.026559

**Authors:** Charlotte Kaplan, Sam J. Kenny, Xuyan Chen, Johannes Schöneberg, Ewa Sitarska, Alba Diz-Muñoz, Matthew Akamatsu, Ke Xu, David G. Drubin

**Affiliations:** Department of Molecular and Cell Biology, University of California, Berkeley, CA 94720-3220; Department of Chemistry, University of California, Berkeley, CA 94720-3220; Center for Neural Circuits and Behavior, University of California, San Diego, CA 92093; Cell Biology and Biophysics Unit, European Molecular Biology Laboratory Heidelberg, Meyerhofstrasse 1, 69117 Heidelberg; Chan Zuckerberg Biohub, San Francisco, CA 94158

**Author notes:** Correspondence to DGD (MA or KX. Collaboration for joint PhD degree between EMBL and Heidelberg University, Faculty of Biosciences. **Preprint Server:** This manuscript is on bioRxiv: https://www.biorxiv.org/content/10.1101/2020.04.05.026559. Classification: Cell biology. Author contributions: C. Kaplan and D.G. Drubin conceived the study and experiments. C. Kaplan performed live cell data acquisition, SRM data analysis and live cell data analysis. S. J. Kenny, S. Chen and K. Xu performed SRM, super-resolution data reconstruction and supervised SRM imaging. J. Schöneberg supported the SRM data analysis. E. M. Sitarska and A. Diz-Muñoz performed membrane tether pulling experiments by atomic force microscopy, data analysis and supervised AFM tether pulling experiments. C. Kaplan, and M. Akamatsu prepared the plot layouts and figures. C. Kaplan, M. Akamatsu, and D.G. Drubin wrote the manuscript with feedback from the other authors.

## Abstract

Clathrin-mediated endocytosis (CME) robustness under elevated membrane tension is maintained by actin assembly-mediated force generation. However, whether more actin assembles at endocytic sites in response to increased load, as has been observed in lamellipodia, has not previously been investigated. Here actin network ultrastructure at CME sites was examined under low and high membrane tension. Actin and N-WASP spatial organization indicate that actin polymerization initiates at the base of clathrin-coated pits and that the network then grows away from the plasma membrane. Actin network height at individual CME sites was not coupled to coat shape, raising the possibility that local differences in mechanical load feedback on assembly. By manipulating membrane tension and Arp2/3 complex activity we tested the hypothesis that actin assembly at CME sites increases in response to elevated load. Indeed, in response to elevated membrane tension, actin grew higher, resulting in greater coverage of the clathrin coat, and CME slowed. When membrane tension was elevated and the Arp2/3 complex was inhibited, shallow clathrin-coated pits accumulated, indicating that this adaptive mechanism is especially crucial for coat curvature generation. We propose that actin assembly increases in response to increased load to ensure CME robustness over a range of plasma membrane tensions.

## Introduction

Actin networks produce force for a wide variety of cellular processes through a Brownian ratchet mechanism (Mogilner and Oster, 1996, 2003; Pollard, 2016). Live-cell studies of lamellipodia (Mueller *et al*., 2017), biochemical reconstitutions (Bieling *et al*., 2016; Funk *et al*., 2021; Li *et al*., 2021) and modeling studies (Akamatsu *et al*., 2020) have shown that actin networks nucleated by the Arp2/3 complex respond to increased load by becoming more dense, which enhances force production. This phenomenon has been demonstrated in the context of actin networks that generate pushing forces. However, actin networks can also generate pulling forces, for example during vesicle formation, but whether networks in this context show load adaptation has not been investigated.

From yeast to humans, transient actin assembly is associated with the formation of clathrin-coated endocytic vesicles. In yeast cells, actin assembly is required to generate forces to invaginate the plasma membrane against a high intrinsic turgor pressure for CME (Kaksonen *et al*., 2005; Aghamohammadzadeh and Ayscough, 2009; Idrissi *et al*., 2012; Kukulski *et al*., 2012). When actin assembly is perturbed in mammalian cells, CME typically slows in a manner that depends on cell type (Fujimoto *et al*., 2000; Merrifield *et al*., 2005; Yarar *et al*., 2005, 2007; Grassart *et al*., 2014; Dambournet *et al*., 2018; Schöneberg *et al*., 2018). A potential cause of this reported variation between cell types might be differences in plasma membrane tension (Pontes *et al*., 2017; Djakbarova *et al*., 2021). Indeed, in mammalian cells, actin assembly becomes increasingly critical as plasma membrane tension increases (Yarar *et al*., 2005; Liu *et al*., 2009; Batchelder and Yarar, 2010; Boulant *et al*., 2011). Actin perturbation results in the accumulation of “U-shaped” membrane invaginations, reflecting difficulty in progressing to the subsequent “Ω-shaped” membrane stage (Fujimoto *et al*., 2000; Yarar *et al*., 2005; Boulant *et al*., 2011; Almeida-Souza *et al*., 2018). In total, these findings suggest that actin assembly improves the efficiency of CME in mammalian cells, potentially compensating for changes in plasma membrane tension.

Despite the fact that actin assembly appears to be associated with CME in all eukaryotes and over a large range of membrane tension values, whether it adapts to changes in membrane tension is not known. Knowing whether the actin cytoskeleton at CME sites adapts to changes in membrane tension will facilitate elucidation of the fundamental mechanisms by which cytoskeletal complexes produce force.

Platinum replica electron microscopy of cultured cells led to the proposal that actin networks assemble in a collar-like arrangement around the vesicle neck (Collins *et al*., 2011). This actin organization would imply that a constricting force is generated toward the neck of the pit, supporting fission. However, actin filaments interact not only with the vesicle neck, but also with the bud surface, given that the clathrin coat is impregnated with actin-binding linker proteins like Hip1R and Epsin (Engqvist-Goldstein *et al*., 2001; Messa *et al*., 2014; Sochacki *et al*., 2017; Clarke and Royle, 2018). Such an arrangement and photobleaching studies imply that actin filaments also apply a force that pulls the clathrin-coated pit into the cell interior (Akamatsu *et al*., 2020).

In yeast, such a pulling mechanism is likely. Actin filaments are nucleated in a ring surrounding the pit and grow from the membrane toward the cell interior, and the resulting filaments are coupled to the clathrin coat surface, generating an inward force orthogonal to the plane of the plasma membrane (Kaksonen *et al*., 2003; Carroll *et al*., 2012; Skruzny *et al*., 2012, 2015; Picco *et al*., 2015; Mund *et al*., 2018). Because the endocytic machinery is highly conserved from yeast to mammals, a similar mechanism for actin force generation during CME seems likely (Akamatsu *et al*., 2020). However, ultrastructural evidence for actin organization through different stages of mammalian CME, and for how this organization might respond to changing membrane tension, are lacking.

Several competing models for actin organization at CME sites in mammalian cells have been proposed, so it is important to distinguish between these models (Engqvist-Goldstein *et al*., 2001; Boulant *et al*., 2011; Messa *et al*., 2014; Sochacki *et al*., 2017; Clarke and Royle, 2018). Recent advances in super-resolution microscopy (SRM) permit examination of cellular ultrastructure with large sample sizes, low invasiveness and high molecular specificity to reveal the ultrastructure of membrane cytoskeletal systems in mammalian cells (Hauser et al., 2018; Xu et al., 2012, 2013).

Here, we combined two-color, three-dimensional stochastic optical reconstruction microscopy (2c-3D STORM) and live-cell fluorescence imaging to determine how filamentous actin is organized at CME sites with different coat geometries under varying values of plasma membrane tension. These measurements lead to the conclusion that the actin network grows from the base of the pit inward, supporting a pulling mechanism for mammalian endocytic actin filaments. The size of the actin network is not tightly coupled to the geometry of the endocytic coat, implying that assembly is modulated by factors uncoupled from coat shape. Importantly, under elevated membrane tension, the actin network grows higher relative to the pit at all endocytic coat geometries. These observations support a mechanism in which the actin network adapts to load by increasing in size to enhance force production and ensure robust CME across a wide range of values of membrane tension.

## Results

### Actin organization at CME sites suggests that force generation can be both parallel and orthogonal to the axis of clathrin-coated pit formation at different sites in the same cell

We used 2c-3D STORM to determine the ultrastructural organization of the actin cytoskeleton at sites of clathrin-mediated endocytosis. Henceforth, we refer to clathrin-coated structures (CCSs) as relatively flat clathrin structures, and clathrin-coated pits (CCPs) as curved, invaginating clathrin structures. Our 2c-3D STORM method preserves cellular ultrastructure by chemical fixation of intact cells, provides high molecular specificity due to immunofluorescence labeling, and allows large numbers of sites to be imaged (Xu *et al*., 2012). We conducted our experiments on a skin-melanoma cell line (SK-MEL-2) wherein ∼87% of dynamin2-eGFP^EN^ (DNM2-eGFP^EN^) spots co-accumulate with actin (Grassart *et al*., 2014). We used this SK-MEL-2 cell line endogenously expressing DNM2-eGFP^EN^ and clathrin light chain A-tagRFP-T (CLTA-TagRFP-T^EN^) for both live-cell and super-resolution experiments (Doyon *et al*., 2011).

In these cells, we resolved CF680-labeled CCSs in the X-Y plane as discrete, round or elliptical shapes on the ventral surface (Fig. 1A). The majority of the CCSs appeared connected to filamentous actin visualized by Alexa Fluor 647-tagged phalloidin. These super-resolution reconstructions resolve the association between clathrin coats and actin networks for hundreds of pits with high resolution in all 3 dimensions. The standard deviations of positions of single fluorophores were 10 nm in-plane for XY and 19 nm in-depth for the Z dimension (Figure S1, Material and Methods).

**Figure 1:**
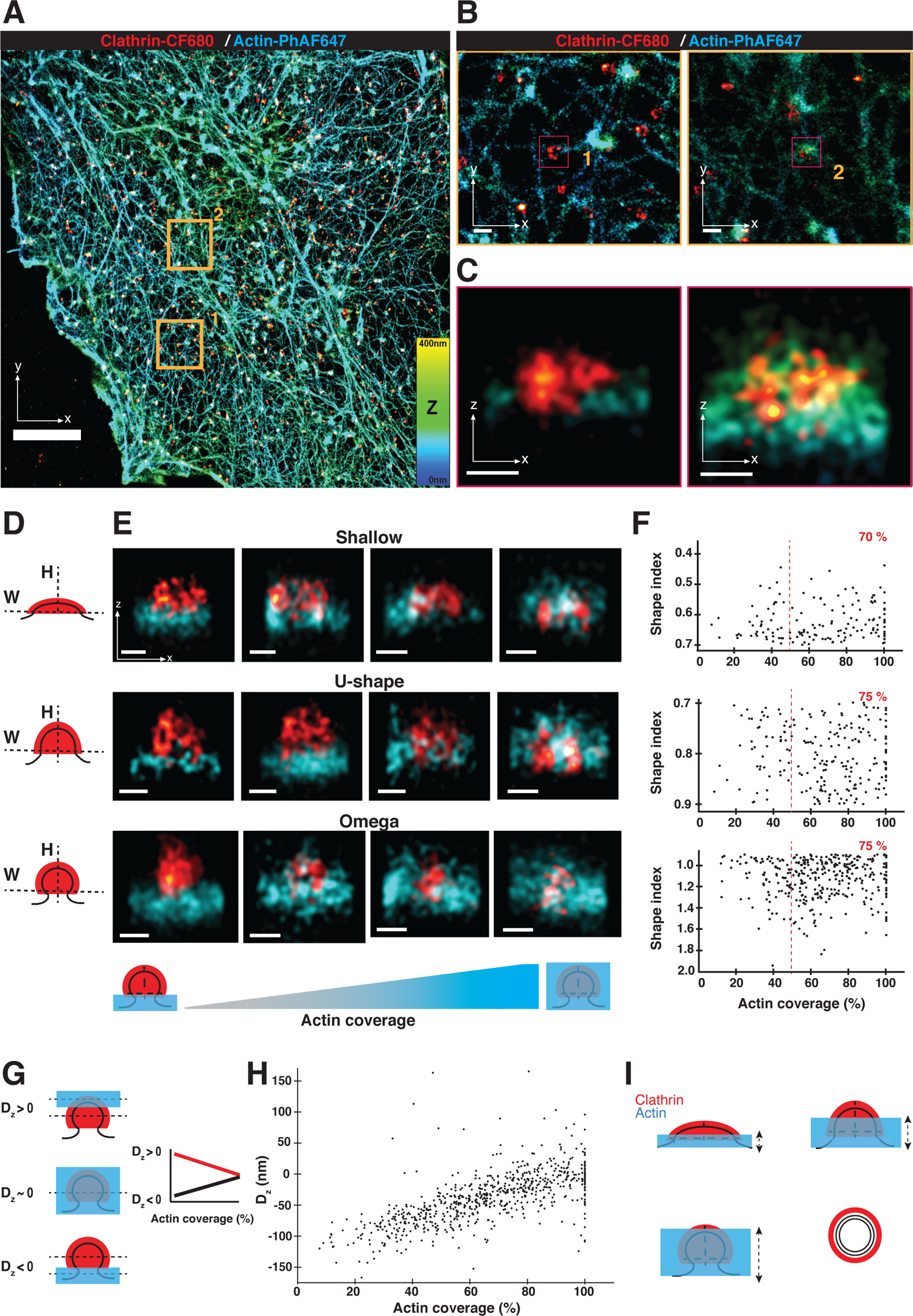
Two-color, three-dimensional stochastic optical reconstruction microscopy (2c-3D STORM) resolves clathrin structures highly connected to actin networks at different stages of endocytosis. **(A)** STORM image of the ventral surface of an SK-MEL-2 cell immunolabeled with the CF-680 antibody (clathrin coats in red) and phalloidin-AF647 (actin in cyan). Orange squares are areas shown in panel B. Color bar shows the z position of actin. Scale bar: 5 µm. **(B)** Magnification of highlighted areas 1 and 2 in panel A. Magenta squares are shown in panel C. Scale bars: 250 nm. **(C)** X-Z projections of the regions highlighted in panel B. Scale bars: 100 nm. **(D**) Illustration of binning clathrin coats (red) into three geometric stages based on their aspect ratio (shape index SI). Shallow: SI < 0.7; U-shape: 0.7 > SI < 0.9 and omega: SI>0.9. **(E)** X-Z projections of representative STORM images showing clathrin coats (red) with different actin (cyan) coverages around clathrin. Calculated shape index of shallow CCSs from left to right image: 0.56, 0.53, 0.51, 0.55; for U-Shaped CCPs from left image to right image: 0.87, 0.89, 0.86, 0.82; for omega-shaped CCPs from left image to right image: 1.31, 1.06, 1.31, 1.52. Scale bars: 100 nm. (**F)** Graphs of endocytic coat shape index as a function of actin coverage for shallow, U-shaped and omega-shaped pits. Pits with actin coverage >5% are shown. Values in red are the percentages of clathrin coats with high (>50%) actin coverage. Upper plot R= − 0.11, n = 150; middle plot R = 0.05, n = 220, lower plot R = − 0.01, n = 347. Events accumulated from 6 cells. **(G)** Cartoon depicting the clathrin coat with actin either at the tip of the coat (top), covering the clathrin coat completely (middle), or at the base of the clathrin coat (bottom). Dashed black lines indicate the average Z position of actin and clathrin. D_z_ is the difference between average actin and clathrin Z positions. D_z_ < 0 is defined as the average actin Z position nearer the base of the pit. Schematic is a hypothetical plot of D_z_ versus actin coverage for scenarios in which actin grows from the tip of the coat (red line) or the base of the pit (black line). **(H)** D_z_ as a function of actin coverage (for actin coverage >5%, R=0.66, n = 719, N_cells = 6). **(I)** Cartoon of actin (blue) growing from the base of the pit (black lines) to cover clathrin coat (red) from a shallow membrane invagination to a fully formed membrane vesicle. X-Z projection (side profile) is shown. Dashed arrows indicate that growth of the actin network is not tightly coupled to the endocytic coat geometry and is variable in extent.

Knowing how actin networks are organized spatially in three dimensions at CME sites provides insights into its force generation mechanisms. It was important to show that we could distinguish actin specifically associated with CCSs from actin in the cell cortex. In Figure S2, we show STORM images to compare actin at CCSs with actin at randomly selected regions of the cell cortex. We found examples of actin that specifically accumulates at the CCP (Figure S2 D and I). Here, actin builds up higher into the cells compared to the actin extending horizontally away from the CCP and at randomly selected sites on the cell cortex (Figure S2 E and J). This observation is consistent with recent cryo-electron tomograms in this cell type distinguishing branched actin networks around clathrin-coated pits from the largely unbranched cortical actin network (Serwas *et al*., 2021).

To then investigate actin organization at multiple CCSs, we rendered the CCSs in three dimensions by cropping a tight area around each clathrin and actin structure in x-y STORM image projection and generated an x-z STORM image projection from this selected ROI (Fig. 1B and C). To our surprise, we observed strikingly different actin filament spatial organizations in the clathrin coats we examined, even when they were near each other in the same cell and even when they were at the same morphological stage of CME (see below) (Fig. 1C). In the first example shown, a thin layer of actin filaments resided along the base of the clathrin coat (Fig. 1C, inset 1), reminiscent of structures observed in EM by Collins *et al*. (Collins *et al*., 2011). In contrast, in the second example, actin filaments covered the CCP completely (Fig. 1C, inset 2, Figure S2 D). This organization resembles actin interacting with the entire clathrin coat as in yeast (Mulholland, 1994; Idrissi *et al*., 2008; Ferguson *et al*., 2009; Kukulski *et al*., 2012; Buser and Drubin, 2013; Mund *et al*., 2018) and in recent cryo-electron tomograms (Akamatsu *et al*., 2020). These micrographs indicate that distinct CME-associated actin structures can coexist in the same cell, consistent with models for force generation both parallel to and orthogonal to the invagination axis.

### CME site-associated actin networks grow from the CCP base to the tip of the coat and their organization is not coupled to clathrin coat geometry

We used quantitative analysis of these STORM reconstructions to determine how actin is organized at CME sites and how actin organization relates to coat geometry. We first selected 989 high-resolution clathrin coats by visual inspection based on quality control criteria explained in the Materials and Methods and determined which ones had associated actin filaments. When cropping CCSs for analysis, we could clearly distinguish single sites from double CCSs. In Figure S3 we present views of 20 sites identified as singles and 20 sites identified as doubles in this manner. Besides the double CCSs, we also excluded CCSs of non-circular shape, as well as very large and very small CCSs in the XY plane (Figure S4 A). We detected actin associated with 74% of the clathrin coats, which is comparable to previous measurements for the same cell type that we made of endocytic traces in live cells of dynamin2-GFP events associated with actin-RFP, given that our super-resolved images are snapshots of a time-lapse event (Grassart *et al*., 2014).

We classified the coats by stage based on their shape in the XZ dimension, similar to earlier analyses of electron micrographs (Avinoam *et al*., 2015). For presentation purposes, clathrin coat shapes were binned into three geometrical categories based on their aspect ratio (which we call the shape index): shallow (flatter coat), U-shaped (intermediate aspect ratio), or Ω-shaped (coat with height similar to its width) (Figs. 1D, S4B and C). This measurement of shape index was consistent whether we carried out the measurement on XZ or YZ projections of the coat (r^2^ = 0.92) (Figure S5). We found a surprisingly wide variety of actin organizations in each of the three geometries (Fig. 1E and S4D). To quantify the relationship between actin organization and endocytic coat geometry, the shape index of the coat was plotted as a function of the extent of actin covering clathrin. When the actin network is larger in size than the clathrin coat, we define this state as 100% coverage (see Materials and Methods, Fig. S4B). There was no significant correlation between actin/coat coverage and coat shape for any of the three geometries (Fig. 1F). We also observed a lateral mean displacement between the peak signals of clathrin and actin of 74 ± 42 nm, indicating an asymmetry in actin localization around the pits (Yarar *et al*., 2005; Collins *et al*., 2011; Jin *et al*., 2021; Serwas *et al*., 2021). This asymmetry value did not significantly change between CME geometries (Fig. S4E and F). We conclude that irrespective of coat geometry, some pits have a thin actin network at the base of the pit, others have an intermediate level of coverage around the clathrin coat, and others have actin completely covering the pit. Thus, actin assembly at CCSs is not coupled to a particular stage of coat curvature development, but might be recruited to promote curvature development.

In our reconstructions, we observed that whenever actin only partially covers the clathrin coat, the network is always located at the base of the pit (Fig. 1E). This observation suggests that actin polymerization is nucleated at the base of the pit and that the network then grows inward around and over the tip of the clathrin coat. To test the generality of this observation, we calculated the difference in average axial (Z) position between the clathrin and actin signals for each site. We define this difference, D_z_, such that negative D_z_ corresponds to actin residing nearer the base of the pit (Fig. 1G). To determine whether actin grows from the base or tip of the pit, we plotted D_z_ as a function of the extent of coverage between actin and the clathrin coat. Increasing values of D_z_ would indicate that the network grows from the base of the pit toward the cell interior (Fig. 1G). Indeed, as a function of actin/coat coverage, D_z_ increased from negative values to near zero (Fig. 1H). We conclude that actin polymerization is initiated at the base of clathrin coats.

Given our finding that actin growth originates from the base of the pit, we next investigated the spatial distribution of the actin nucleation factor N-WASP at clathrin coats by 2c-3D STORM. Consistent with our conclusions about where actin assembly occurs at CCPs, N-WASP localized to the base of both shallow and highly curved clathrin coats (Fig. S6A). More unexpectedly, at some CME sites, N-WASP covered the entire clathrin coat irrespective of coat geometry (Fig. S6B).

In summary, we conclude that actin polymerization is nucleated at the base of clathrin-coated pits and grows toward the coat’s tip (Fig. 1I). Unexpectedly, actin nucleation is not coupled to coat geometry. A possible explanation for the variety of actin organizations we observed associated with clathrin coats is that actin network organization responds to differences in load at CME sites.

### Dynamics of clathrin-mediated endocytosis slow down under elevated membrane tension

We next combined osmotic shock with live-cell fluorescence microscopy and 2c-3D STORM to determine how actin-mediated force generation contributes to clathrin-coated pit formation under elevated membrane tension (Fig. 2A). Previous EM studies identified a requirement for actin filaments at the “U” to “Ω” transition (Boulant *et al*., 2011). However, for a mechanistic understanding, the quantitative relationship between membrane tension and endocytic dynamics must be elucidated (Akamatsu *et al*., 2020). Our quantitative light microscopy-based analysis of a large number of sites at different CME stages provided the necessary sensitivity to detect effects throughout the process. We first needed to establish conditions under which CME dynamics are affected by elevated membrane tension in these cells. To determine how membrane tension is affected by changes in media osmolarity, we performed membrane tether pulling experiments by atomic force microscopy (AFM) on SK-MEL-2 cells cultured under isotonic conditions and hypotonic conditions (75 mOsm) (Fig. 2B). In isotonic media, the force required to maintain a pulled membrane tether was 33.0 ± 7.4 pN. Under hypotonic conditions the tether force increased to 48.0 ± 17.1 pN (Fig. 2B). These measurements allowed us to quantitatively relate a given hypotonic environment in these cells to changes in membrane tension.

**Figure 2:**
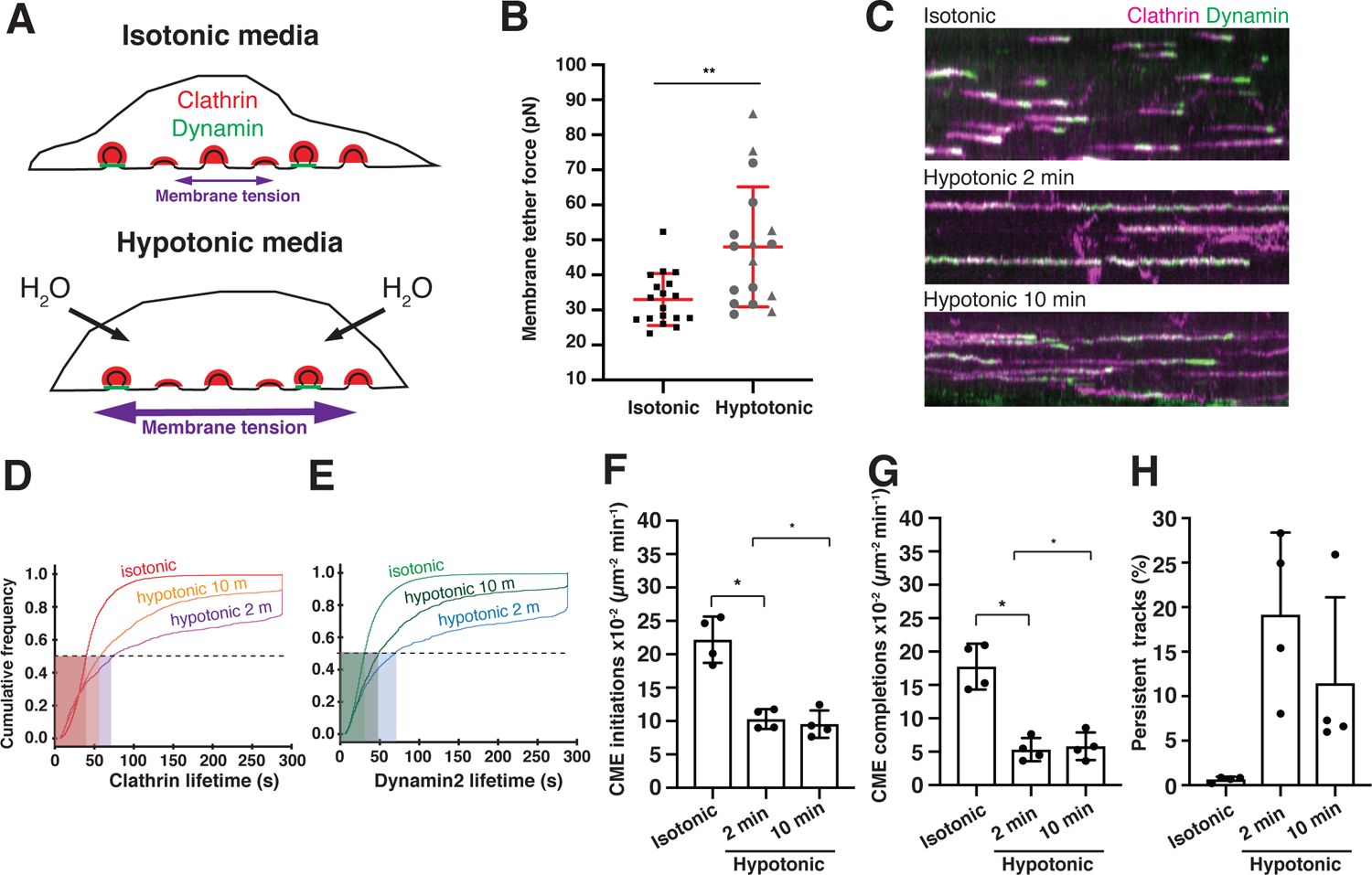
Quantitative analysis of clathrin-mediated endocytosis mechanosensitivity under elevated membrane tension. **(A)** Schematic of cells in isotonic media (top) or hypotonic media (bottom), which causes water influx and stretches the cell membrane. In this figure, hypotonic treatment is 75 mOsm. **(B)** Mean membrane tether force values measured by atomic-force microscopy of cells in isotonic media (n = 18) or in hypotonic media (n = 17). Mean values were obtained by pulling at least 3 tethers (3 independent experiments). In hypotonic treatment, circles are mean tether values from 2 min to 10 min after hypotonic media exchange, and triangles are mean tether values obtained between 10 min and 16 min after hypotonic media exchange. Bars are mean ± SD. p = 0.002 by Mann-Whitney test. **(C)** Kymographs of total-internal reflection fluorescence micrographs of live SK-MEL-2 cells endogenously expressing CLTA-TagRFP-T^EN^ (magenta) and DNM2-eGFP^EN^ (green). Time is on the X axis. Kymographs are 4.8 min long. Cells were imaged in isotonic media (top), or hypotonic media for 2 min (middle) or 10 min (bottom). **(D)** Cumulative distribution plot of clathrin lifetimes marked by CLTA-TagRFP-T^EN^ in isotonic media (red), hypotonic media for 2 min (violet), and hypotonic media for 10 min (orange). These tracks were associated with DNM2-eGFP^EN^. **(E)** Cumulative distribution plot of dynamin2 lifetime marked by DNM2-eGFP^EN^ in isotonic media (light green), hypotonic media for 2 min (blue), and hypotonic media for 10 min (dark green). These tracks were associated with CLTA-TagRFP-T^EN^. n = (5831) tracks in 17 cells across four experiments for D-H. **(D)** and **(E)** detailed statistics in Table S1. **(F)** Plot of endocytic initiation rate for the three conditions. p<0.05 by Kolmogorov-Smirnov test for both comparisons. **(G)** Endocytic completion rate in the 3 conditions. p<0.05 by Kolmogorov-Smirnov test for both comparisons. **(H)** Percentage of persistent tracks (defined as tracks lasting the entirety of the image acquisition) for the three conditions. **(F)** - **(H)** Boxplots show mean ± SD. Statistics in Table S1 and Fig. S7H.

To decipher the relationship between CME dynamics and membrane tension, we used total internal reflection fluorescence (TIRF) microscopy to image SK-MEL-2 cells under isotonic and hypotonic conditions (Fig. 2C). CLTA-TagRFP-T^EN^ and DNM2-eGFP^EN^ fluorescence lifetimes were determined by single-particle tracking (Hong *et al*., 2015). In isotonic media most clathrin tracks were relatively short (47 ± 32 s) with a burst of DNM2-eGFP^EN^ signal peaking at the end (DNM2 lifetime 39 ± 32 s) (Fig. 2C, Supp. movie 1, Table S1). Neither isotonic media nor exchange to slightly hypotonic 225 mOsm media noticeably affected DNM2-eGFP^EN^ and CLTA-TagRFP-T^EN^ lifetimes or CME initiation and completion rates (Fig. S7 A-D). Only 1 - 2% of these CLTA-TagRFP-T^EN^ and DNM2-eGFP^EN^ fluorescence tracks persisted over the entire 4.8 min movie under these conditions (Fig. S7 B and D).

Upon treatment with moderately hypotonic (150 mOsm) media for 10 min, CLTA-TagRFP-T^EN^ lifetime in cells increased by 20% (62 ± 51 s versus 49 ± 31 s) (Fig. S7E). This moderate treatment also had mild effects on the CME initiation rate (15 ± 5 x 10^-2^ µm^-2^ min^-1^ versus 25 ± 6 x 10^-2^ µm^-2^ min^-1^), completion rate (11 ± 4 x 10^-2^ µm^-2^ min^-1^ versus 19 ± 6 x 10^-2^ µm^-2^ min^-1^), and stalling rate (events that persist the entire length of our 4.8 min movies) (3 ± 1 % versus 0.8 ± 0.9 % (Fig. S7F).

Treatment of cells with more strongly hypotonic (75 mOsm) media dramatically perturbed CME dynamics. After 2 minutes in 75 mOsm hypotonic media, tracks elongated and very often lasted the entire duration of the 4.8 min movie (Fig. 2C, Supp. movie 2, Table S1). The mean lifetimes of CLTA-TagRFP-T^EN^ and DNM2-eGFP^EN^ tracks were 128 ± 112 s (Fig. 2D) and 125 ± 112 s, respectively (Fig. 2E). We also observed a substantial decrease in CME initiation rate (10.3 ± 1.5 x 10^-2^ µm^-2^ min^-1^ versus 22.2 ± 3.5 x 10^-2^ µm^-2^ min^-1^), completion rate 5.3 ± 1.7 x 10^-2^ µm^-2^ min^-1^ versus 17.8 ± 3.4 x 10^-2^ µm^-2^ min^-1^), and an increase in track stalling (19 ± 9 % versus 0.7 ± 0.3 %) (Fig. 2F-H, Fig. S7H). After 10 min of culturing, the CLTA-TagRFP-T^EN^ and DNM2-eGFP^EN^ lifetimes began to recover, most likely reflecting cellular adaptation to the hypotonic treatment (Fig. 2C-H, Supp. movie 3). We did not detect effects of hypotonic media treatment on lifetimes of tracks containing only CLTA-TagRFP-T^EN^, characteristic of clathrin structures not associated with CME (Fig. S7A,C,E,G) (Hong *et al*., 2015). DNM2-eGFP^EN^-only events showed a moderate response to elevated membrane tension. We conclude that elevating plasma membrane tension with hypotonic shock markedly perturbs CME dynamics in a dose-dependent manner.

### Actin force generation assists clathrin coat remodeling under elevated membrane tension

Next, we determined the role of branched actin filament assembly on endocytic outcome under elevated membrane tension. Branched actin networks are generated by activated Arp2/3 complex (Mullins *et al*., 1998). We inhibited Arp2/3 complex-mediated actin polymerization using the small molecule CK666 (Nolen *et al*., 2009; Hetrick *et al*., 2013). We reasoned that inhibition of the Arp2/3 complex might have less effect on membrane tension than global actin inhibitors such as latrunculin and jasplakinolide often used in previous CME studies. Nevertheless, since Arp2/3 complex inhibition can affect membrane tension (Diz-Muñoz *et al*., 2016), we first carefully established experimental conditions in which effects of high membrane tension or CK666 would not mask one another. Treatment of cells with 100 µM CK666 did not affect the membrane tether force in these cells (p>0.5) during the time the experiment was performed (Fig. S8A). Under these optimized conditions, we performed live-cell fluorescence microscopy and STORM to learn more about the clathrin coat remodeling steps at which branched actin network assembly is required.

We first titrated CK666 and monitored the effect on CME dynamics after 2 min of treatment. We aimed to identify a minimal CK666 concentration that induced a rapid effect on CME dynamics. 100 µM CK666 extended lifetimes of CLTA-TagRFP-T^EN^ to 79 ± 66 s after 2 mins of treatment compared to 56 ± 40 s for the DMSO control (Figs. S8B, Table S2). Similarly, DNM2-EGFP^EN^ lifetimes increased to 65 ± 64 s upon CK666 treatment, compared to 49 ± 38 s for the DMSO control (Figs. S8B, Table S2). 100 µM CK666 did not affect CME completion frequency or the percentage of persistent CLTA-TagRFP-T^EN^ tracks (Fig. S8 C and D), though we observed a small decrease in the CME initiation rate (Fig. S8E).

The elongation of DNM2-eGFP^EN^ and CLTA-TagRFP-T^EN^ lifetimes upon 100 µM CK666 treatment was exacerbated upon simultaneous incubation in moderately hypotonic media (150 mOsm). Compared to controls, the combination of 100 µM CK666 and 150 mOsm hyopotonic media markedly lengthened the clathrin lifetimes to 96 ± 86 s (compared to DMSO control 59 ± 52 s) and dynamin2 to 84 ± 85 s (compared to DMSO control 47 ± 49 s) (Fig. 3A-F, Table S3). We conclude that in these cells, Arp2/3 complex-mediated actin assembly is required to maintain normal CME dynamics under elevated membrane tension.

**Figure 3:**
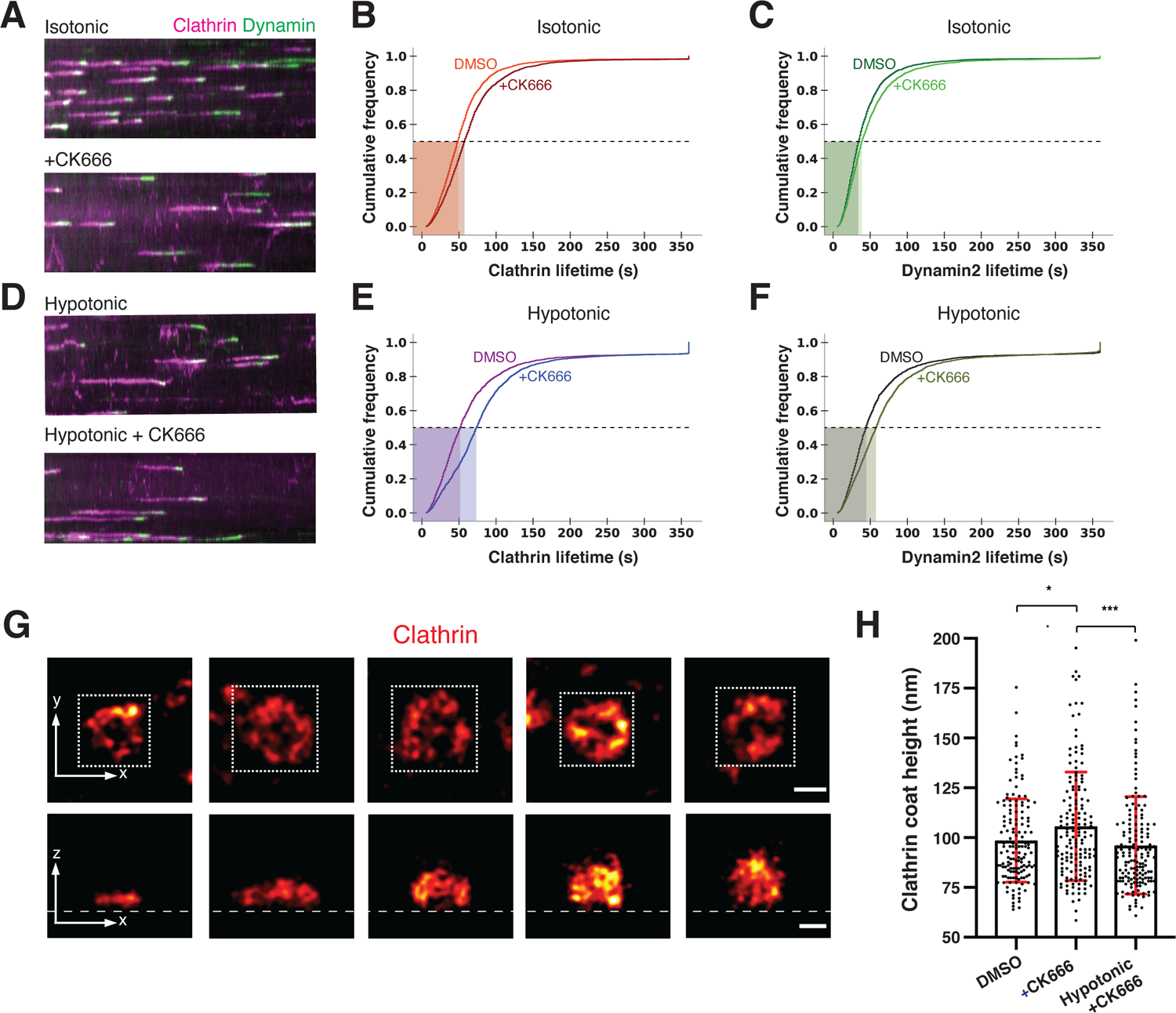
Importance of Arp2/3 complex-mediated actin polymerization during CME increases under elevated membrane tension. In this figure, CK666 (Arp2/3 complex inhibitor) treatment is 100 µM and hypotonic shock is 150mOsm. **(A)** Kymographs of cells expressing CLTA-TagRFP-T^EN^ (magenta) and DNM2-eGFP^EN^ (green) imaged by total-internal reflection microscopy. Cells were imaged in isotonic media. The media was then exchanged to include CK666 (lower panel) and imaged after 4 min. **(B)** Cumulative distribution plots of clathrin lifetimes in control, DMSO-treated (orange, (n = 4219)), and CK666-treated (red, n = 3124) conditions. **(C)** Cumulative distribution plots for control, DMSO-treated (dark green, n = 4219), and CK666-treated (light green, n = 3124) dynamin2 lifetimes associated with clathrin lifetimes in (B). (B) and (C) control N_cells = 10 and CK666 treatment N_cells = 10 measured in 3 independent experiments. Complete statistics in Table S3. **(D)** Kymographs of cells in hypotonic media. In the upper panel, cells were placed in hypotonic media and imaged after 4 min. In the lower panel, cells were treated with CK666 in hypotonic media and imaged after 4 min. **(E)** Cumulative distribution plots of clathrin lifetimes for control, DMSO-treated (magenta, n = 1405), and CK666-treated (blue, n = 2783) in hypotonic media. **(F)** Cumulative distribution plots of DMSO-treated (black, n = 1405) and CK666-treated (olive, n = 2783) dynamin2 lifetimes in hypotonic media associated with clathrin lifetimes in (E). **(E)** and **(F)** control N_cells = 9 and CK666 treatment N_cells = 10 measured in 3 independent experiments. Complete statistics in Table S3. **(G)** Representative STORM images of immunolabeled clathrin-coated structures in control cells arranged by coat height. Upper panel shows the x-y projections and the lower panel the corresponding x-z projections. The white square in the x-y projections shows the area that was cropped to generate the x-z projections. The height of clathrin coats in the x-y projection from left to right image is 61 nm, 96 nm, 124 nm, 158 nm and 177 nm. Scale bars: 100 nm. **(H)** Clathrin coat heights when cells were treated with DMSO (n = 154) or CK666 (n = 158) in isotonic media or CK666 in hypotonic media (n = 159). Clathrin coat images for quantitative analysis were collected from at least 3 cells for each condition. Statistics are given in Figure S8F. p<0.05 in both comparisons by Mann-Whitney test.

Next, we used STORM to identify which endocytic coat geometries were enriched upon a combination of Arp2/3 complex inhibition and osmolarity treatment. Cells were cultured under the CK666 and osmolarity conditions described above, chemically fixed, and then immunolabeled for clathrin. Clathrin coat height served as a metric for the membrane invagination stage. As in the above 2d-3D STORM experiments, the full progression from a flat clathrin coat to a rounded vesicle could be clearly resolved in 3D (Fig. 3G, lower image panel). Control cells treated with DMSO showed an average clathrin coat height of 98 ± 21 nm (Fig. 3H, Fig. S8 F). The average height increased to 106 ± 27 nm when cells were treated with 100µM CK666 (Fig. 3J). Thus, when Arp2/3 complex activity is inhibited in these cells, more clathrin pits accumulate at a greater height. This suggests that more clathrin pits stall at a later stage of progression, consistent with observations of accumulated coat geometries from actin inhibitor studies (Yarar *et al*., 2005; Boulant *et al*., 2011; Yoshida *et al*., 2018).

Interestingly, when Arp2/3-mediated actin polymerization was inhibited in cells with elevated membrane tension, the average clathrin coat height decreased to 96 nm ± 24 nm (Fig. 3G and H, Fig. S8F). This height decrease was also reflected in an accumulation of smaller shape indices and a very mild effect on clathrin coat width (Fig. S8 F – H). This result, along with our observation of slowed CME dynamics, suggests that upon membrane tension elevation in response to hypotonic conditions in SK-MEL-2 cells, inhibition of Arp2/3 complex activity may cause an enrichment of shallow endocytic coat geometries.

### Actin organization adapts to elevated membrane tension by increasing clathrin coat coverage

Finally, we used STORM to determine the relationship between elevated membrane tension and actin cytoskeleton organization at CME sites (Fig. 4). We treated cells with strong (75 mOsm) hypotonic shock for 5 min and then chemically fixed them for 2c-3D STORM. When we super-resolved clathrin and actin by 2c-3D STORM in these cells, the actin cytoskeleton remained intact (Pan *et al*., 2019) and associated with CCSs (Fig. S9 A - D).

**Figure 4:**
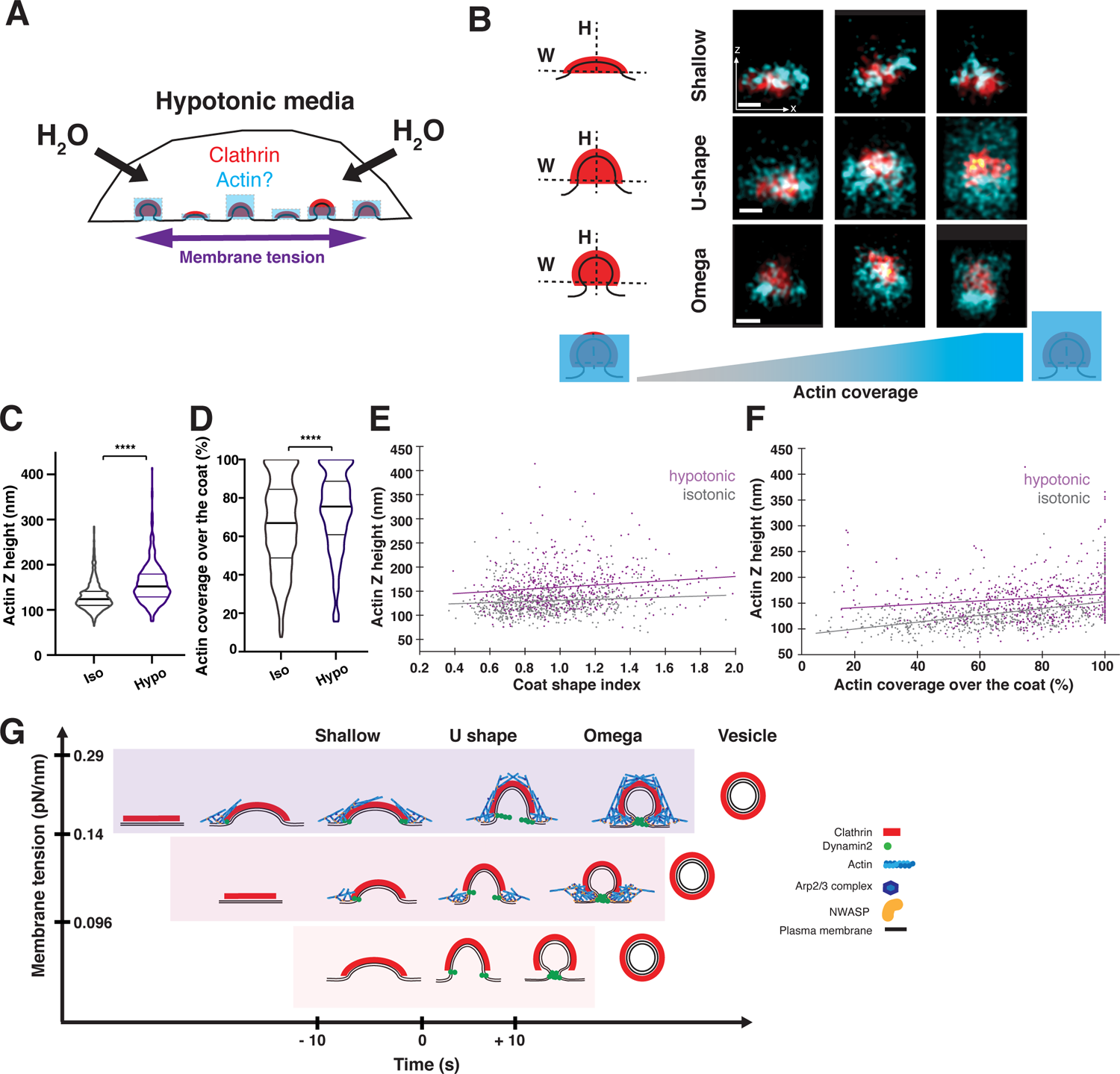
The actin network at CME sites increases in size in response to elevated membrane tension. In this figure, hypotonic refers to 75 mOsm media. **(A)** Schematic of cells in hypotonic media, which increases plasma membrane tension. The response of the actin network (blue) to elevated plasma membrane tension (purple) was previously unknown. **(B)** Representative STORM images of clathrin (red) and actin (cyan) in x-z projections for cells fixed after treatment in the hypotonic media for 5 min (bottom). Coated pits are classified as shallow, U-shaped, or omega-shaped based on the aspect ratio of the coat. Scale bars: 100 nm. **(C)** Plots of actin Z height at clathrin-coated pits from cells in the isotonic (n = 736) and hypotonic (n = 527) media measured from STORM x-z projections. Lines are median ± interquartile range. p<0.0001 determined by Mann-Whitney test. **(D)** Plots of actin coverage over the clathrin coat in pits found in STORM x-z projection images in isotonic (n = 719) and hypotonic (n = 509) conditions. Pits with actin coverage > 1% are plotted. Lines are median ± interquartile range. p<0.0001 determined by Mann-Whitney test. **(E)** Actin Z height as a function of coat shape in isotonic (gray, n = 736) and hypotonic (purple, n = 527) conditions. **(F)** Actin Z height as a function of actin coverage over the clathrin coat in isotonic (gray, n = 719) and hypotonic (purple, n = 509) conditions. The data for isotonic conditions were also used to generate the plots in Figure 1. Three independent STORM experiments with N_cells = 6 in isotonic and N_cells = 7 in hypotonic media. **(G)** Cartoon depicting an adaptive actin force-generating mechanism that counteracts elevated membrane tension to ensure robust CME progression. This schematic describes three scenarios in which membrane tension is low, intermediate or high, and how CME is accomplished efficiently by adaptive actin network organization. Under low tension (bottom), the clathrin coat provides sufficient force to advance CME. At intermediate tension (middle), actin polymerization is required for the transition from U to omega shape. At high tension (top), endocytic progression slows. More pits stall at the shallow conformation. In response to increased resistance, the actin network grows to envelop the coat and provide additional force against high membrane tension.

Strikingly, in response to hypotonic media treatment, the average actin height dramatically increased for all endocytic geometries (Fig. 4B, C and S9 E-G). When we examined randomly selected regions of the cell cortex, we did not observe any increase in cortical actin height, providing evidence that any observed effects would be specific for CME sites (Figure S10). Overall, the average actin height at CME sites increased from 130 ± 30 nm under isotonic media conditions to 160 ± 40 nm (Fig. 4C). This increase corresponded to an increase of actin growth over the clathrin coat from covering 66 ± 23% of the coat to covering 73 ± 21% under 75 mOsm hypotonic media (Fig. 4D). The increase in actin height was observed for all coat shapes (e.g., shallow or curved pits) (Fig. 4E). Similarly, for different extents of clathrin coverage by actin, the average actin height was greater in hypotonic conditions across different levels of coat coverage (Fig. 4F). However, the extent of asymmetry between actin and clathrin signals did not significantly change compared to the isotonic condition (Fig. S9G). These observations of greater average actin height and coverage over the clathrin coat suggest that the force contribution of actin at CME sites is increased when membrane tension is elevated such that a greater load is carried by the actin network.

Overall, these observations lead us to conclude that under elevated plasma membrane tension, actin grows higher in the z-dimension around clathrin coats (Fig. 4G). Such an adaptive mechanism for actin organization presumably generates the required forces to ensure the efficient progression of mammalian CME under varying levels of membrane tension.

## Discussion

By combining two-color, three-dimensional STORM imaging and quantitative live-cell TIRF microscopy for thousands of CME sites, with AFM tether-pulling membrane tension measurements, and manipulations of membrane tension and Arp2/3-mediated actin assembly, we showed that actin assembly adapts to changes in load at individual CME sites. This mechanism likely ensures that sufficient forces are generated for robust endocytic progression over a range of membrane tension regimes. While STORM of individual CCSs cannot attain the resolution of EM, our approach had several advantages that allowed us to gain new mechanistic insights: (1) it allowed us to sample much larger numbers of CME sites with a preserved actin network than is possible by EM, thus permitting rigorous quantitative analysis, (2) we imaged CME sites and associated actin in intact cells that had not been subjected to unroofing or extraction protocols, and (3) we were able to use antibodies and fluorescent phalloidin to unambiguously identify specific proteins at CME sites.

### Actin assembly and organization adapt to elevated membrane tension

Elevating membrane tension can have a dramatic impact on CME progression in mammalian cells (Raucher and Sheetz, 1999; Boulant *et al*., 2011; Ferguson *et al*., 2016, 2017; Willy *et al*., 2017; Bucher *et al*., 2018). However, how cells adapt to compensate for increased load at CME sites has not been elucidated. Our results provide critical mechanistic insights into how the CME machinery adapts to elevated membrane tension to maintain robust CME progression. We showed quantitatively that actin assembly and organization adapt to changes in membrane tension, which we measured by AFM membrane tether pulling (Fig. 2). Changes in membrane tension can in principle occur globally (entire cells) or locally (in different regions of one cell, or even within different regions of a single endocytic site) (Houk *et al*., 2012; Rangamani *et al*., 2014; Shi *et al*., 2018). Since we detect different actin organizations at individual CME sites with similar clathrin coat geometries within a single cell, such differences might reflect subcellular, local load variation resulting from variance in such factors as membrane tension and cell adhesion. Observation in other studies that actin assembles late in CME progression (Merrifield *et al*., 2002; Grassart *et al*., 2014; Akamatsu *et al*., 2020; Jin *et al*., 2021), coupled with the observation here that actin can associate with clathrin coats of different geometries, is consistent with the possibility that clathrin lattices of low curvature can develop high curvature under the “constant area” model (Avinoam *et al*., 2015; Sochacki *et al*., 2021). Moreover, our data provide evidence that the low to high coat curvature transition becomes more dependent on actin assembly forces as load increases.

### Measurement of membrane tension changes using atomic force microscopy

Using atomic force microscopy, we measured the membrane tether force for SK-MEL-2 cells in isotonic and hypotonic conditions (Fig. 2). We measured tension on the dorsal cell surface while imaging endocytic events on the ventral surface. Local differences in membrane tension have been observed in some cell types (Shi *et al*., 2018)but not others (Houk *et al*., 2012). Nevertheless, AFM remains the gold standard for measuring apparent membrane tension in cells (Sitarska and Diz-Muñoz, 2020). In isotonic conditions, the tether force was 33 ± 7 pN (Fig. 2B). This tether force is within an intermediate range measured for other cell types such as NIH3T3 cells, HeLa cells, and macrophages (Sitarska and Diz-Muñoz, 2020; Roffay *et al*., 2021). Assuming a 100 pN*nm bending rigidity of the plasma membrane, this value corresponds to a membrane tension of 0.14 ± 0.04 pN/nm (Diz-Muñoz *et al*., 2013). In hypotonic conditions, the membrane tether force increased to 48 ± 17 pN, which corresponds to a doubling of membrane tension to 0.29 ± 0.04 pN/nm. Higher membrane tether values have been reported for other cell types (Sitarska and Diz-Muñoz, 2020). Below, we describe implications of earlier results and our new results reported here on actin’s role in CME under three different plasma membrane tension regimes (Fig. 4G).

### Low membrane tension regime

When membrane tension is low, clathrin coat assembly provides sufficient energy to bend the underlying plasma membrane into a full spherical shape, similar to the constant curvature model (Fig. 4G) (Saleem *et al*., 2015; Willy *et al*., 2021). We indeed found in our STORM data that 26% of clathrin coats lack actin in shallow, U-shape and omega-shaped CME geometries. This observation is consistent with mathematical modeling, which indicates that the coat can provide sufficient energy to bend the plasma membrane when membrane tension is low (0.002 pN/nm) (Hassinger *et al*., 2017). Here, we consider low membrane tension to be of a value lower than those we measured for SK-MEL-2 cells in standard media (Fig. 2). Given that actin polymerization appears to be dispensable for CME in some cell types, even though it still might make CME more efficient, we hypothesize that the basal membrane tension may be lower in those cell types. However, we caution the interpretation of experiments with harsher actin drug treatments, as prolonged actin inhibitor treatment of cells can perturb the actin cortex and reduce the tether force measurements (membrane tension) by a factor of ∼50% (Pontes *et al*., 2017). In future studies, strategies should be developed to relate local membrane tension to specific endocytic site geometry and actin organization.

### Intermediate tension regime

In the intermediate tension regime, defined as the resting tension of SK-MEL-2 cells in isotonic conditions (Fig. 2), clathrin coat assembly and membrane curvature-inducing proteins still appear to provide sufficient energy to drive clathrin coats to adopt the U shape (Fig. 4, intermediate membrane tension). When we inhibited Arp2/3-mediated actin polymerization using CK666, clathrin coat progression stalled at an aspect ratio most likely reflecting the U-shaped stage, consistent with the effects of actin assembly inhibition reported for other cell types (Yarar *et al*., 2005; Boulant *et al*., 2011; Almeida-Souza *et al*., 2018). Thus, at intermediate membrane tension it appears that actin force generation is primarily required for the U-shaped to Ω-shaped clathrin coat transition. The actin network observed near the base of the pit may reflect a role driving plasma membrane neck constriction and scission by generating forces orthogonal to the direction of membrane invagination as proposed previously (Bucher *et al*., 2018; Scott *et al*., 2018; Mund *et al*., 2021).

### High membrane tension regime

Our live-cell and STORM observations indicate that as membrane tension is elevated further, coat deformation and membrane invagination during the early stages of CME become increasingly dependent upon actin force generation (Fig. 4). When membrane tension was elevated to an intermediate level (e.g., 150 mOsm media), CME lifetimes slowed modestly (Fig. S7). When we inhibited Arp2/3-mediated actin polymerization using CK666 in cells treated with 150 mOsm media, the average clathrin coat height was lower than in CK666-treated cells cultured under isotonic conditions, likely reflecting an enrichment of shallow pits (Fig. 3H). Under elevated membrane tension, the actin cortex tends to become thinner and less dense (Houk *et al*., 2012; Chugh *et al*., 2017; Roffay *et al*., 2021). However, our study shows that the actin network surrounding clathrin-coated pits instead increases in height under elevated membrane tension (Figure 4). Actin assembly from the base of the CCP continues until the network covers the clathrin coat completely, allowing it to interact with proteins linking the actin network to the clathrin coat (Engqvist-Goldstein *et al*., 2001; Messa *et al*., 2014; Sochacki *et al*., 2017). Actin-binding linker proteins such as Hip1R and Epsin1 cover the clathrin coat completely and are thus positioned to provide high internalization efficiency by harnessing actin assembly forces perpendicular to the plasma membrane (Akamatsu *et al*., 2020; Serwas *et al*., 2021). It may be that the “constant area” model (Avinoam *et al*., 2015; Sochacki *et al*., 2021), in which a flat or shallow coat grows to full size and then bends, applies under the high load regime (Bucher et al., 2018; Mund et al., 2021; Scott et al., 2018), and it is here where actin assembly forces are most important for remodeling the coat into a curved vesicle.

An important question for future studies concerns the nature of the adaptive mechanism that increases actin assembly in response to elevated membrane tension. Possible mechanisms include: (1) stalling works passively to allow actin to assemble longer; or (2) high load works actively through mechanisms documented for other contexts, for example by increasing contact of filaments associated with the coated pit with membrane-associated N-WASP-Arp2/3 complex, leading to more assembly, alongside load-dependent decreases in the rate of filament capping (Akamatsu et al., 2020; Bieling et al., 2016; Funk et al., 2021; Li et al., 2021). As endocytosis and other actin-mediated trafficking events operate within a different membrane geometry and local protein and lipid environment than in lamellipodia, a high priority is now to determine whether similar adaptive mechanisms operate during such local membrane bending processes.

When the actin network fully covers the clathrin coat, it resembles the radial organization described by mathematical modeling for mammalian CME and the actin network organization described for budding yeast (Ferguson *et al*., 2009; Hassinger *et al*., 2017; Mund *et al*., 2018; Akamatsu *et al*., 2020). In yeast this actin organization drives endocytic membrane invagination against the high resistance that results from turgor pressure. Mathematical modeling showed that this organization produces high forces perpendicular to the plasma membrane (Hassinger *et al*., 2017). Actin-generated forces parallel and orthogonal to the membrane invagination at high tension may coexist to drive membrane invagination followed by neck constriction and scission.

CME dynamics dramatically slow down when cells are in this high membrane tension regime, resulting in only ∼40% of endocytic lifetimes shorter than 50 s compared to 80% at low membrane tension (Fig. 2D). We also detected ∼19% of lifetimes being longer than the 4.8 min movies we captured, a 20-fold increase in stalled (persistent) tracks (Fig. 2H). We suggest that this tension regime pushes this adaptive mechanism to the limit.

### N-WASP spatial organization suggests an actin force generation control mechanism

N-WASP spatial organization at CCSs and CCPs provides valuable mechanistic insight into how actin network assembly contributes to force generation during CME. We found that N-WASP localizes at the base of early and late clathrin-coated pits, where it likely interacts with SH3 domain-containing proteins present at the endocytic membrane neck neck (Almeida-Souza et al., 2018; Schöneberg et al., 2017; Sochacki et al., 2017). This organization is similar to that of the homologous nucleation promoting factor Las17 in budding yeast (Mund *et al*., 2018). Filaments nucleated at the base of CCPs would be able to interact with coat proteins such as Hip1R and Epsin1/2/3 to generate forces to invaginate the plasma membrane (Hassinger *et al*., 2017; Mund *et al*., 2018; Akamatsu *et al*., 2020; Joseph *et al*., 2020).

Intriguingly, we also sometimes observed a strikingly different N-WASP spatial organization in which it was distributed over the full clathrin coat. The type II nucleation factors Abp1 and cortactin bind to actin filaments and to the Arp2/3 complex and could serve as binding partners for N-WASP around the clathrin-coated pit (Guo et al., 2018; Helgeson and Nolen, 2013; Le Clainche et al., 2007; Pinyol et al., 2007). This WASP location might reflect a distinct path for filament nucleation, originating from the coat rather than the base, that is potentially important to generate higher forces when actin already surrounds the clathrin-coated pit.

A late burst of actin assembly was shown previously to often accompany CME (Merrifield *et al*., 2002) and to facilitate CME progression, especially when membrane tension is high (Fujimoto *et al*., 2000; Yarar *et al*., 2005; Batchelder and Yarar, 2010; Boulant *et al*., 2011; Grassart *et al*., 2014; Li *et al*., 2015; Ferguson *et al*., 2017; Yoshida *et al*., 2018; Akamatsu *et al*., 2020; Jin *et al*., 2021). In this study we observed that load-adapted, CME-associated actin assembly buffers changes in plasma membrane tension, thereby ensuring clathrin-coated vesicle formation over a range of membrane tension regimes. We expect that the ability of actin assembly to respond to load is a common critical feature in membrane remodeling events.

## Materials and Methods

### Cell culture

SK-MEL-2 cells from clone Ti13 (hCLTA^EN-1^ /hDNM2^EN-1^) were cultured in DMEM/F12 with GlutaMax™ supplement (10565-018, Thermo Fisher Scientific) media containing 10% fetal bovine serum (FBS) and 1,000 U/mL penicillin-streptomycin mix (15140122, Thermo Fisher Scientific) and kept in a 37°C humidified incubator with 5% CO2 (cell source information in (Doyon *et al*., 2011)). After each cell vial was thawed, cells were checked after 2 passages for mycoplasma contamination. Cell line authentication was performed by short tandem repeat validation.

### Antibodies and reagents

The primary antibodies used were mouse anti-clathrin light chain (AB CON.1, MA5-11860, Thermo Fisher Scientific), mouse anti-clathrin heavy chain (AB X-22, MA1-065, Thermo Fisher Scientific) and rabbit anti-N-WASP (ab126626, Abcam). The secondary antibodies used were Alexa Fluorophore 647 chicken anti-rabbit (A21443, Thermo Fischer Scientific), goat anti-mouse (115-005-205, Jackson ImmunoResearch) conjugated to CF680-NHS ester (Biotium 92139). Reagents and small molecule inhibitors used were DMSO (D2650, Sigma Aldrich), CK666 (SML0006, batch # 0000012761, Sigma Aldrich) and Phalloidin-AF647 (A22287, Fisher Scientific).

### Preparation of CF680-labeled secondary goat anti-mouse antibody

CF680 NHS ester was dissolved at a concentration of 3 mM in anhydrous DMSO. 1 µL of dye solution, 80 µL of a 1.25 mg/mL suspension of unlabeled goat anti-mouse IgG1 secondary antibody (115-005-205, Jackson ImmunoResearch Laboratories, Inc.), and 10 µL of 1M sodium bicarbonate solution were mixed and allowed to react for 15 min at room temperature. The reaction mixture was added to an equilibrated NAP-5 column (Sigma GE17-0853-01) and flushed with PBS. The dye conjugated antibody was collected from the first colored eluent fraction and a concentration of 0.12mg/mL was determined with a NanoDrop spectrophotometer.

### Sample preparation for two-color clathrin and actin imaging

18 mm round coverslips were cleaned 20 min in 70% ethanol (Electron Microscopy Science, Cat # 72222-01). Cells were detached with 500uL 0.05% trypsin (25300-054, Gibco), washed once in DMEM/F12 and collected by centrifugation. Cells were counted using a hemocytometer and 20,000 cells/mL were seeded onto 18 mm round coverslips in 12-well plates. Cells were incubated for 16 – 24 hours in culture media prior to preparation for imaging.

Cells were fixed first for 1-2 min in 0.3% (v/v) glutaraldehyde (GA) solution containing 0.25% (v/v) Triton in cytoskeleton buffer (CB: 10mM MES, 150mM NaCl, 5mM EGTA, 5mM Glucose, 5mM MgCl_2_, 0.005% NaN_3_, pH6.1) and then immediately fixed for 10 min in 2% (v/v) GA solution in CB. Both solutions were prepared fresh from a 70% GA stock (Electron Microscopy Science, cat #16365) (protocol follows reference: (Xu *et al*., 2012). After fixation, samples were washed once in CB and then incubated for 7 min in freshly prepared CB containing 0.1% (w/v) NaBH_4_. Subsequently, samples were washed 3 times for 10 min in CB with gentle agitation on a shaker. Samples were then blocked for 30 min in 5% (w/v) BSA in CB (Sigma Aldrich, A3733). For dense clathrin labeling, light (diluted 1:200) and heavy chain (diluted 1:200) antibodies were used together in a 1% (w/v) BSA CB solution. Primary antibody immunostaining was performed overnight at 4°C. On the next day, samples were washed twice in 1% (w/v) BSA CB for 5 min. The mouse secondary antibody-CF680 was used at a final concentration of 0.40 µg/mL – 0.60 µg/mL in a 1% BSA - 1x CB solution. Samples were stained for 30 min at room temperature in the dark and washed twice for 5 min in 1% (w/v) BSA CB solution, and then for 10 min in CB solution. Samples were then placed into a solution of CB containing 0.5µM Phalloidin-AF647 and kept at room temperature in the dark for a minimum of 2 hours. Samples were washed once with PBS before STORM imaging.

### Sample preparation for single-color clathrin and dual-color N-WASP imaging

Cells were prepared as for the two-color sample preparation on coverslips, and then fixed for 20 minutes in 3% (v/v) paraformaldehyde (PFA, 15710 Electron Microscopy Sciences) in CB (protocol follows (Li *et al*., 2018). Samples were washed quickly in CB and subsequently were incubated for 7 min in freshly prepared 0.1% (w/v) NaBH_4_ in CB solution. Subsequently, samples were washed 3 times for 10 min in CB with gentle agitation on a shaker and permeabilized afterwards in a 0.1% Triton-PBS solution for 1-2 min. For single antibody clathrin staining, subsequent washing, blocking and antibody incubation steps were similar to the two-color clathrin and actin sample preparation protocol.

Dual-color immunolabeling was performed with primary antibody against N-WASP (diluted 1:200), clathrin heavy and clathrin light chain (diluted 1:600-1:1000) in 1% (w/v) BSA in PBS overnight at 4°C. Samples were washed the next day twice for 5 min in 1% (w/v) BSA in PBS. Secondary antibody staining was first performed with Alexa Fluorophore 647 anti-rabbit antibody (diluted 1:200) in 1% BSA (w/v) in PBS for 30 min at room temperature and kept in the dark. After two 10 min long washes in PBS containing 1% (w/v) BSA, secondary antibody staining was performed with CF680 anti-mouse antibody (diluted 1:600). The samples were given three final washes in PBS for 10 min each.

### SRM imaging

Dye-labeled cell samples were mounted on glass slides in a standard STORM imaging buffer consisting of 5% (w/v) glucose, 100 mM cysteamine, 0.8 mg/mL glucose oxidase, and 40 µg/mL catalase in 1M Tris-HCL (pH 7.5) (Huang et al., 2008; Rust et al., 2006). Coverslips were sealed using Cytoseal 60. STORM imaging was performed on a homebuilt setup (Wojcik *et al*., 2015) based on a modified Nikon Eclipse Ti-U inverted fluorescence microscope using a Nikon CFI Plan Apo λ 100x oil immersion objective (NA 1.45). Dye molecules were photoswitched to the dark state and imaged using a 647-nm laser (MPB Communications); this laser was passed through an acousto-optic tunable filter and introduced through an optical fiber into the back focal plane of the microscope and onto the sample at an intensity of ∼2 kW cm^2^. A translation stage was used to shift the laser beam toward the edge of the objective so the light reached the sample at incident angles slightly smaller than the critical angle of the glass-water interface. A 405-nm laser was used concurrently with the 647-nm laser to reactivate fluorophores into the emitting state. The power of the 405-nm laser (typical range 0-1 W cm^-2^) was adjusted during image acquisition so that at any given instant, only a small, optically resolvable fraction of the fluorophores in the sample was in the emitting state. For 3D STORM imaging, a cylindrical lens was inserted into the imaging path so that images of single molecules were elongated in opposite directions for molecules on the proximal and distal sides of the focal plane (Huang *et al*., 2008). The raw STORM data were analyzed according to previously described methods (Huang et al., 2008; Rust et al., 2006). Data were collected at a frame rate of 110 Hz for a total of ∼80,000 frames per image. Single and two-color imaging was performed on cells labeled with Alexa Fluor 647 only or Alexa Fluor 647 and CF680 with 647-nm excitation based on a ratiometric detection scheme (Bossi et al., 2008; Gorur et al., 2017; Testa et al., 2010). In the two-color imaging scheme, light emitted from the AF647 and CF680 fluorophores was collected concurrently and split into two light paths using a long pass dichroic mirror (T685lpxr; Chroma). Each light path was projected onto one half of an Andor iXon Ultra 897 EM-CCD camera. Dye assignment was performed by localizing and recording the intensity of each single molecule in each channel. Conventional imaging of 560- and 488-nm dyes was performed immediately prior to STORM imaging using the appropriate laser and filter set. Emission data were collected through the short wavelength reflected path of the aforementioned optical setup and overlaid directly onto the final STORM image. Details of selection and analysis of SRM images are found in the Supporting Information.

Our 3D-STORM setup was based on the same design as (Huang *et al*., 2008) and so we achieved comparable resolutions. The experimental STORM resolution was measured by repeatedly measuring the position of a single fluorophore and determining the standard deviation (SD) of the localization distribution (Huang *et al*., 2008; Xu *et al*., 2015). We accordingly examined our STORM data in this work, and overlayed the localization distributions of 24 representative single molecules from 3 different samples, as shown (Fig. S1). Gaussian fits (red curves) gave standard deviations of 10 nm in-plane for the *XY* directions and 19 nm in-depth for the *Z* direction. These results are similar to those reported in the Figure 1C of (Xu *et al*., 2015), where standard deviations are 9 nm in *X*, 11 nm in *Y*, and 22 nm in *Z*.

### Total internal reflection fluorescence (TIRF) microscopy

TIRF imaging was carried out on a Nikon Eclipse Ti2 inverted microscope with a CFI60 60x Apo TIRF objective and a Hamamatsu Orca-Flash 4.0 V2 sCMOS camera. eGFP and Tag.RFP-T fluorescence were excited using 488 nm and 561 nm lasers and detected using a Chroma HC TIRF Quad Dichroic (C-FL TIRF Ultra Hi S/N 405/488/561/638) and Chroma HC Quad emission filters BP 525/50 and BP600/50, respectively (Bellows Falls, VT). Unless mentioned specifically, channels were acquired sequentially at a 1.2 sec interval and 400ms exposure time over 4.8 minutes to 6 minutes. Real-time acquisition was achieved by a National Instruments (PXI 1033, Austin, TX) controller. The system was controlled with NIS-Elements software and maintained at 37**°**C by an OkoLab environmental chamber (Burlingame, CA).

### Hypo-osmotic media treatment

SK-MEL-2 cells were plated on glass coverslips one day prior to osmotic treatment and imaging: 20,000 cells/ mL were seeded 16h – 24h prior to the experiment on 25 mm round #1.5 glass coverslips that had been cleaned with 70% ethanol (Warner Instruments, 64-0715). Isotonic imaging media contained Dublbecco’s Modified Essential Medium and Ham’s F-12 medium (DMEM/F12) without phenol red (11039, Thermo Fisher Scientific) with 5% v/v FBS. The media was diluted with an inorganic salt solution containing 10mM CaCl_2,_ 0.3mM MgCl_2_ and 0.1mM MgSO_4_ (CMM) to maintain concentrations of critical ions, while obtaining hypo-osmotic conditions by diluting the media containing components such as D-Glucose. 225 mOsm hypotonic imaging media contained 1:4 v/v CMM solution in DMEM/F12, 150 mOsm hypotonic imaging media contained 1:1 v/v CMM solution in DMEM/F12, and 75 mOsm hypotonic imaging media contained 4:1 v/v CMM solution in DMEM/F12. 5% v/v FBS was present in all hypotonic solutions.

### For live cell fluorescence microscopy

CLTA-TagRFP-T^EN^ and DNM2-eGFP^EN^ fluorescence in SK-MEL-2 cells were acquired first in isotonic media over a course of 4.8 minutes. Subsequently, media was exchanged on the stage to hypotonic media (either 225 mOsm, 150 mOsm or 75 mOsm) and movies were acquired for 4.8 minutes, starting 2 minutes and 10 minutes after media exchange. Media exchange on the stage did not affect CME initiation rates or fluorescence lifetimes beyond the existing experimental intrinsic variability (Fig. S7 A,B).

For STORM imaging, 75 mOsm hypotonic buffer treatment was performed in the cell culture dish for 5 min. Cells were immediately chemically fixed after the 5 min treatment and further treated with the STORM sample preparation protocol as described above. Methods to analyze TIRF data can be found in the Supporting Information.

### CK666 concentration titration

20,000 SK-MEL-2 cells/ mL were seeded in 8 well chambers 16h – 24h prior to the experiment (80826, ibidi, Fitchburg, WC). A CK666 (SML0006, batch # 0000012761, Sigma Aldrich) stock solution was prepared at 50mM in DMSO and kept at −20°C. 25µM, 50µM and 100 µM CK666 and equivalent 0.5% v/v DMSO, 1% v/v DMSO and 2% DMSO v/v solutions for controls were prepared fresh in DMEM/F12 containing 5% FBS and kept at 37°C until used. Cells were first imaged in DMEM/F12 containing 5% FBS solution as a baseline control for 4.8 minutes. Subsequently, imaging solution was exchanged on the microscopy stage to CK666 or DMSO containing imaging solution and another 4.8-minute movie was acquired after 2 minutes of treatment. Each treatment was repeated twice and an area of 1024 pixel x 1024 pixel was used to record 3-6 cells per experiment.

### CK666 in combination with hypo-osmotic media

Cells were prepared as for the CK666 concentration titration experiment described above. Solutions of 2% v/v DMSO in DMEM/F12, 100 µM CK666 in DMEM/F12, 2% v/v DMSO in 1:1 v/v CMM solution in DMEM/F12 (150 mOsm hypotonic media) and 100 µM CK666 1:1 v/v CMM solution in DMEM/F12 (150 mOsm hypotonic media) were prepared fresh and kept at 37°C until used. All solutions contained 5% FBS. Cells were first imaged in DMEM/F12-5% FBS solution as a baseline control for 6 minutes. Subsequently, the imaging solution was exchanged on the microscopy stage to the desired experimental solutions and a 6-minute movie was recorded after 4 minutes of incubation.

### Tether pulling experiments using Atomic Force Microscopy

Custom-cut 35-mm glass-bottom dishes (Greiner Bio-One, #627860) were coated with fibronectin (50 ug/mL, Corning #356008) for 30 minutes and washed with DPBS shortly before use. SK-MEL-2 cells were seeded at a density of 0.15-0.20×10^5^ cells/ml in DMEM/F12 GlutaMax^TM^ supplement media with 1% FBS and penicillin-streptomycin mix (Gibco^TM^, #15140-122) in a 37°C humid incubator with 5% CO_2_ for 2-4 hours, and used directly for membrane tether pulling experiments. OBL-10 cantilevers (Bruker) were mounted on a CellHesion 200 AFM (Bruker) integrated into an Eclipse Ti inverted light microscope (Nikon), calibrated using thermal noise method and coated with 2.5 mg/ml Concanavalin A (C5275, Sigma) for 1 hour at 30°C. After rinsing the cantilever with DPBS, it was positioned at any location over the cell for tether pulling using brightfield imaging. Approach velocity was set to 1 µm/s, contact force to 100–300 pN, contact time to 300 ms–10 s, and retraction speed to 10 µm/s. After a 10 µm tether was pulled, the cantilever position was held constant until the moment of tether breakage and at least 2 seconds afterwards. Sampling rate was set to 2000 Hz. After measurements of tether forces in control conditions, an inorganic salt solution containing 10mM CaCl_2,_ 0.3mM MgCl_2_ and 0.1mM MgSO_4_ was added to the medium (4:1 v/v) to achieve 75 mOsm hypotonic treatment. Tether forces were measured after media dilution for 2-16 minutes. Tether forces per cell are the average of at least 3 tethers. Cells were not used longer than 1 h for data acquisition. Force-time curve analysis was performed using the JPKSPM Data Processing Software.

### Data analysis, statistical analysis and data plotting

For statistical analysis and data plotting, Prism version 7.0e and matplotlib in a Jupyter notebook (5.5.0) were used.

## Acknowledgments

CK was funded by the German research foundation (DFG KA4305/1-1). DGD was funded by NIH grant R35GM118149. KX was funded by NSF under CHE-1554717 and the Pew Biomedical Scholars Award. ADM was funded by the European Molecular Biology Laboratory (EMBL), the Human Frontiers Science Program (HFSP) grant number RGY0073/2018 and the Deutsche Forschungsgemeinschaft (DFG) grant numbers DI 2205/2-1 and DI 2205/3-1. MA was funded by NIH grant 1 K99 GM132551-01. ES was funded by the EMBL and the Joachim Herz Stiftung Add-on Fellowship for Interdisciplinary Science. We thank Justin Taraska and Kem Sochacki (NHLBI, NIH) for use of the platinum replica EM images for comparison in Figure S3. We thank Yidi Sun, Padmini Rangamani and Ross T.A. Pedersen for critical reading and discussions on the manuscript. We thank Sungmin Son and Daniel A. Fletcher for valuable input and discussion on the manuscript.

## Abbreviations

CME: Clathrin-mediated endocytosis

CCV: Clathrin coated vesicles

CCS: clathrin coated structure

CCP: Clathrin coated pits

SI: shape index

