## Supplementary figures 1-10 for "Load adaptation of endocytic actin networks"

**Figure S1**

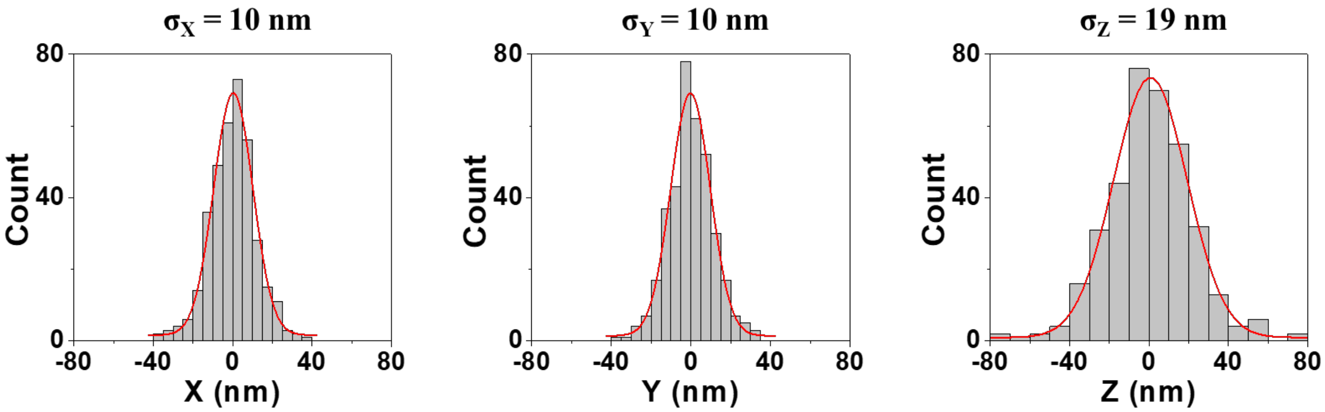

$\sigma_X$ = 10 nm
$\sigma_Y$ = 10 nm
$\sigma_Z$ = 19 nm
Count
80
40
0
-80
-40
0
40
80
X (nm)
Y (nm)
Z (nm)

### Figure S2

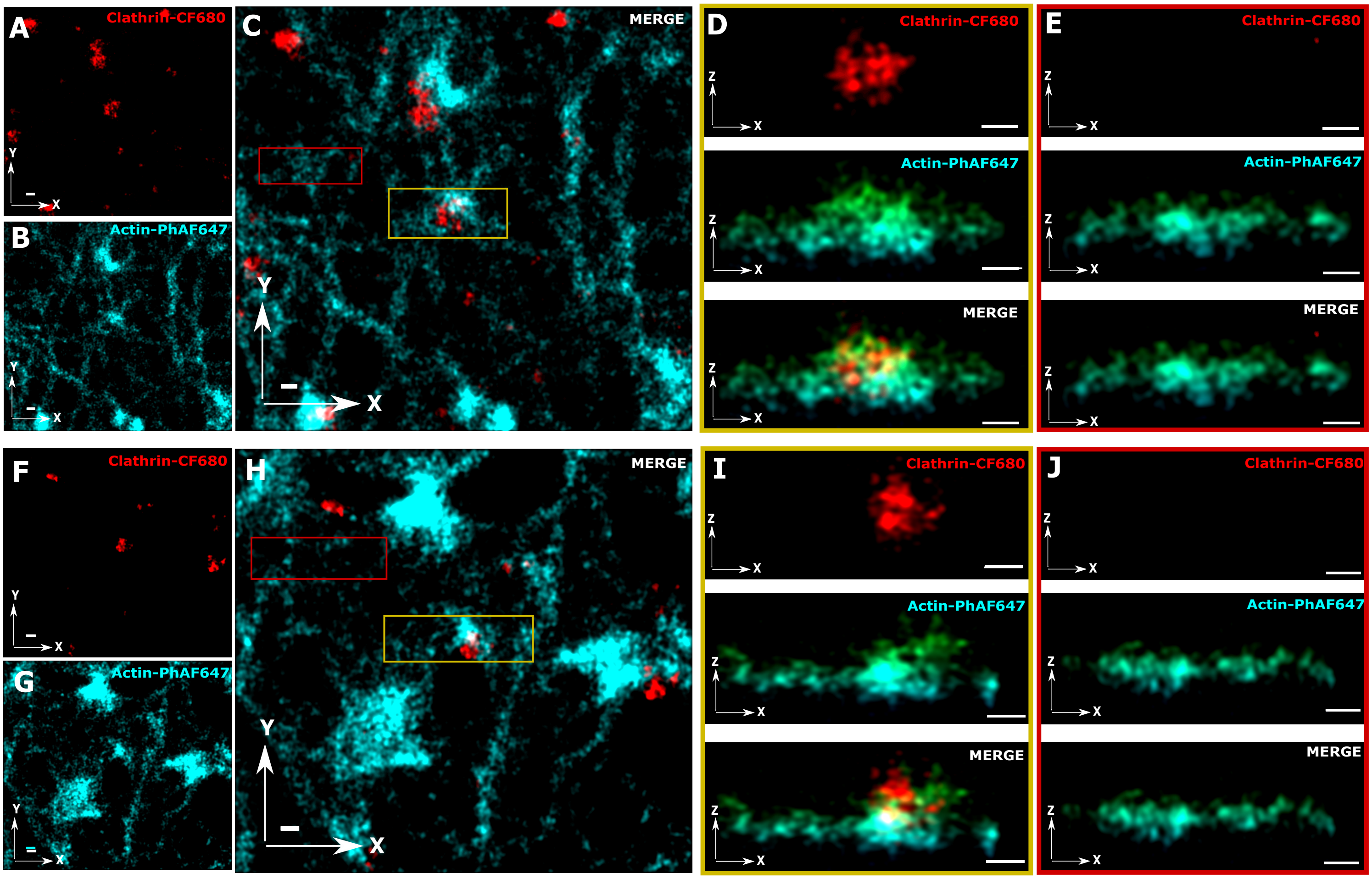

A
Clathrin-CF680
B
Actin-PhAF647
C
MERGE
D
Clathrin-CF680
Actin-PhAF647
MERGE
E
Clathrin-CF680
Actin-PhAF647
MERGE
F
Clathrin-CF680
G
Actin-PhAF647
H
MERGE
I
Clathrin-CF680
Actin-PhAF647
MERGE
J
Clathrin-CF680
Actin-PhAF647
MERGE

Figure S3

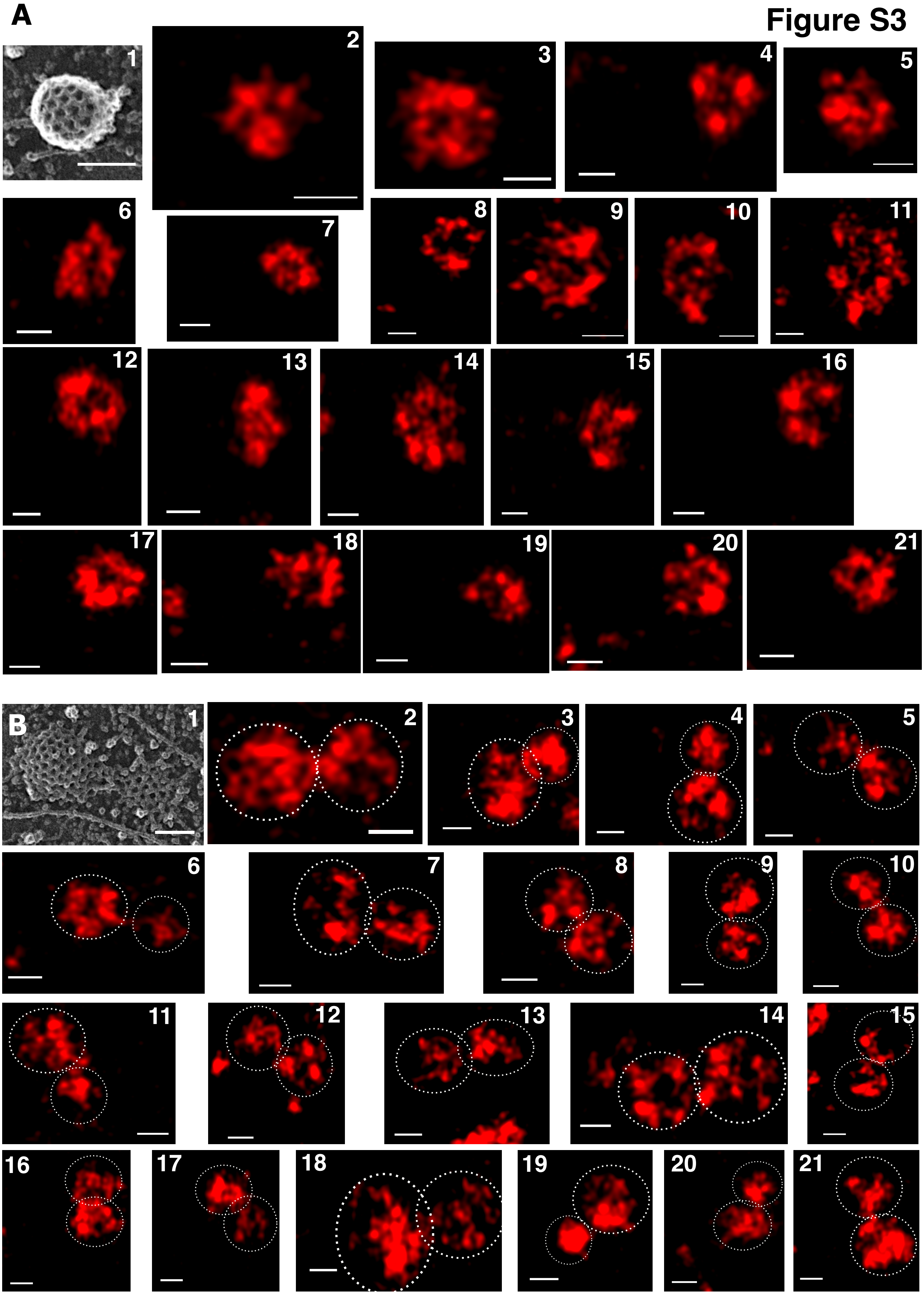
A
B

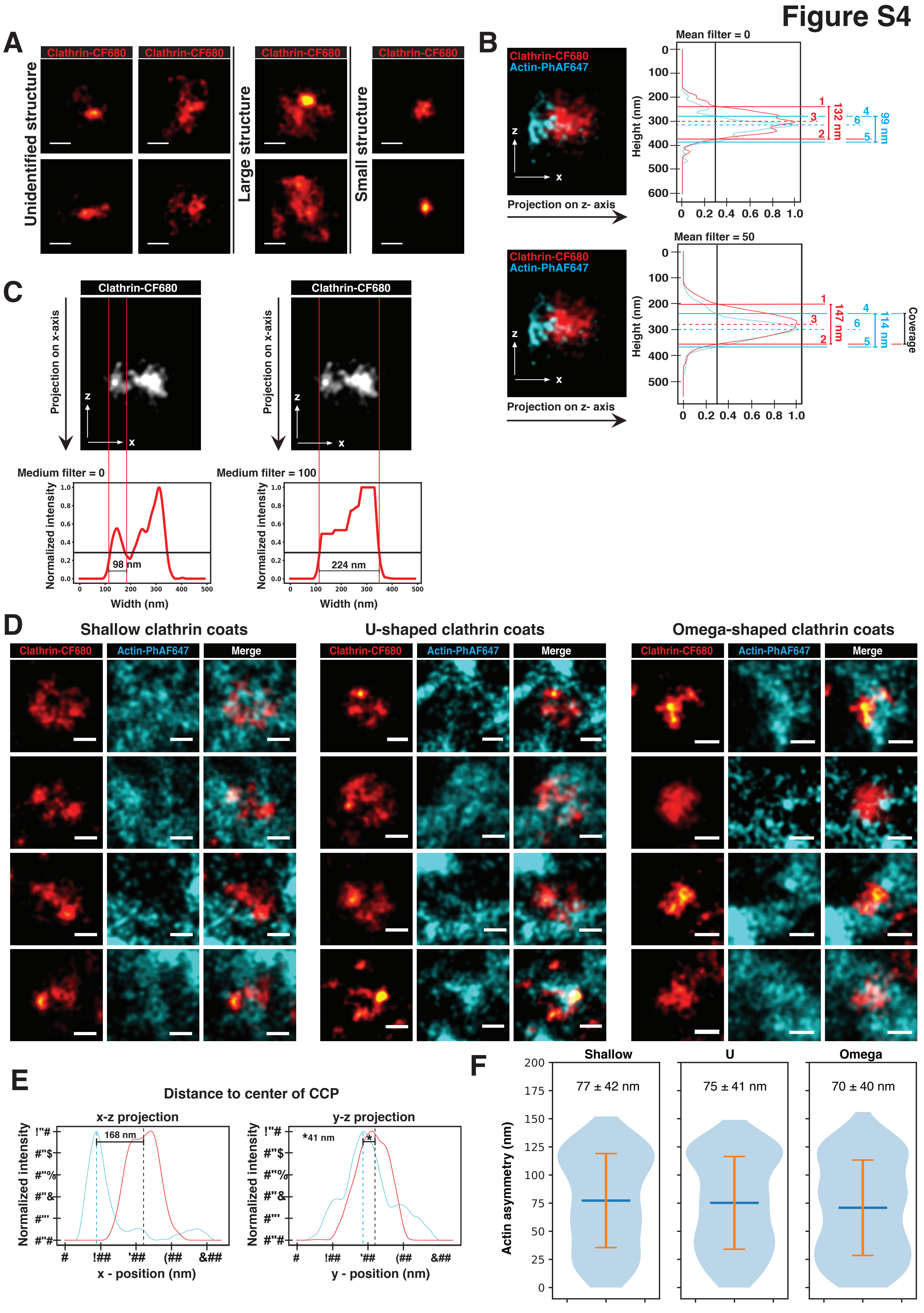

Figure S4
A
Clathrin-CF680
Unidentified structure
Large structure
Small structure
B
Clathrin-CF680
Actin-PhAF647
Projection on z- axis
Mean filter = 0
Height (nm)
132 nm
99 nm
Mean filter = 50
147 nm
114 nm
Coverage
C
Clathrin-CF680
Projection on x-axis
Medium filter = 0
Medium filter = 100
Normalized intensity
98 nm
224 nm
Width (nm)
D
Shallow clathrin coats
U-shaped clathrin coats
Omega-shaped clathrin coats
Clathrin-CF680
Actin-PhAF647
Merge
E
Distance to center of CCP
x-z projection
y-z projection
Normalized intensity
168 nm
*41 nm
x - position (nm)
y - position (nm)
F
Shallow
U
Omega
77 ± 42 nm
75 ± 41 nm
70 ± 40 nm
Actin asymmetry (nm)

### Figure S5

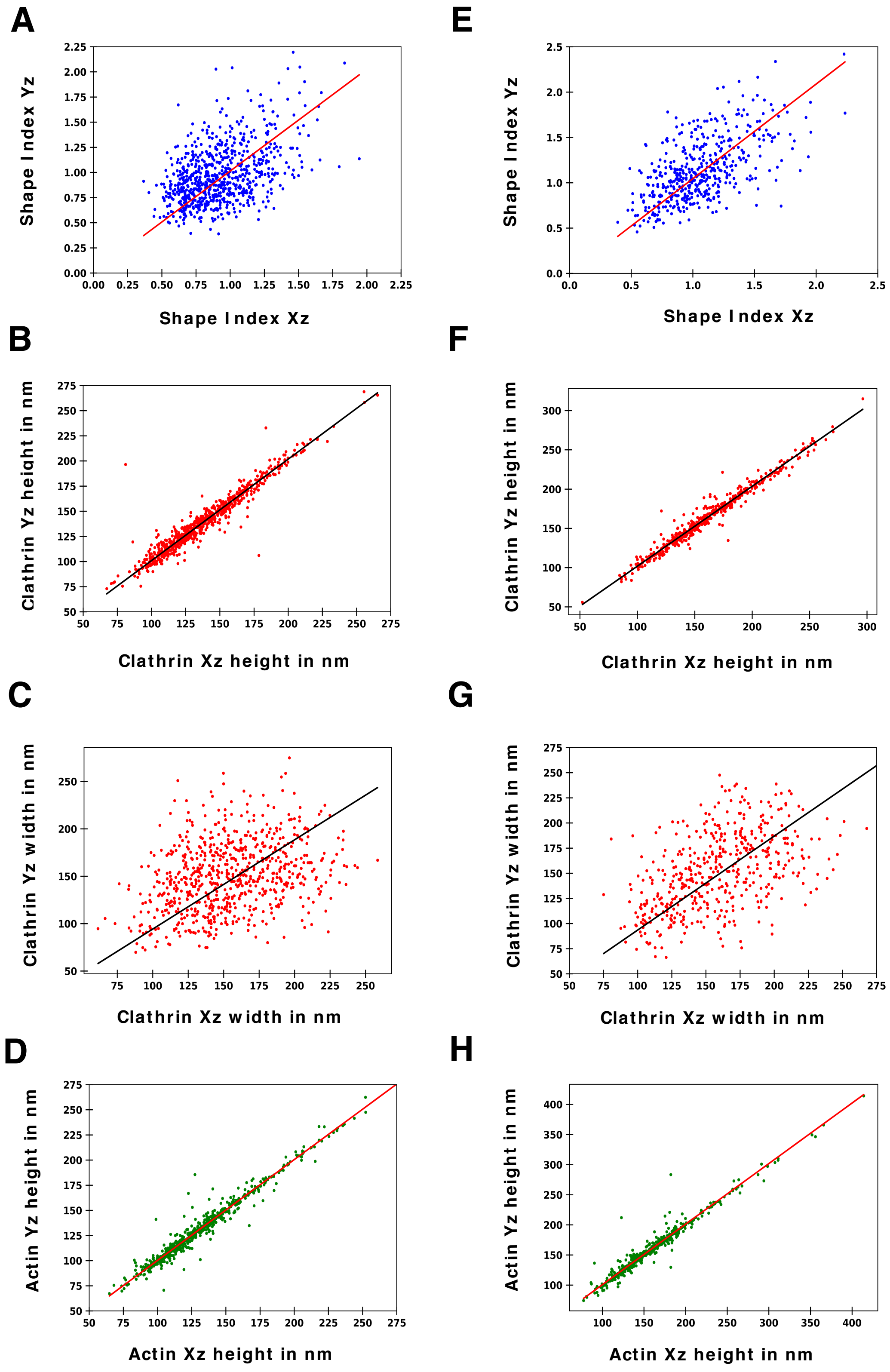
A
Shape Index Yz
Shape Index Xz
E
Shape Index Yz
Shape Index Xz
B
Clathrin Yz height in nm
Clathrin Xz height in nm
F
Clathrin Yz height in nm
Clathrin Xz height in nm
C
Clathrin Yz width in nm
Clathrin Xz width in nm
G
Clathrin Yz width in nm
Clathrin Xz width in nm
D
Actin Yz height in nm
Actin Xz height in nm
H
Actin Yz height in nm
Actin Xz height in nm

### Figure S6

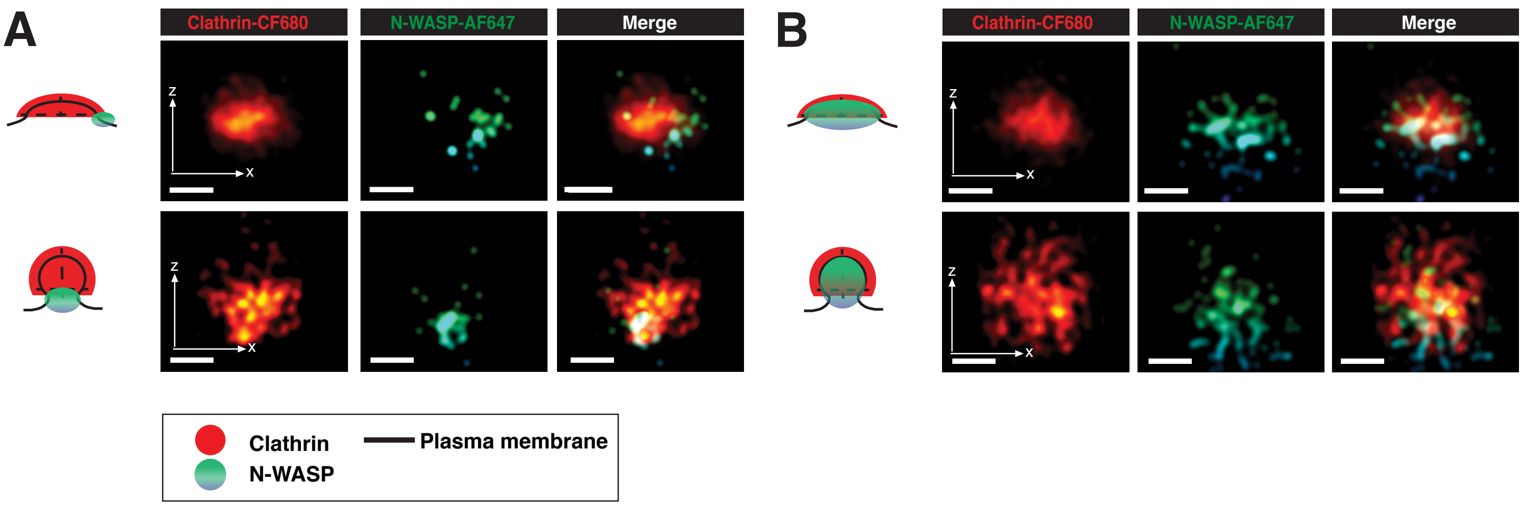
A
Clathrin-CF680
N-WASP-AF647
Merge
z
x
B
Clathrin-CF680
N-WASP-AF647
Merge
Clathrin
N-WASP
Plasma membrane

### Figure S7

**A** Isotonic media, 300 mOsm

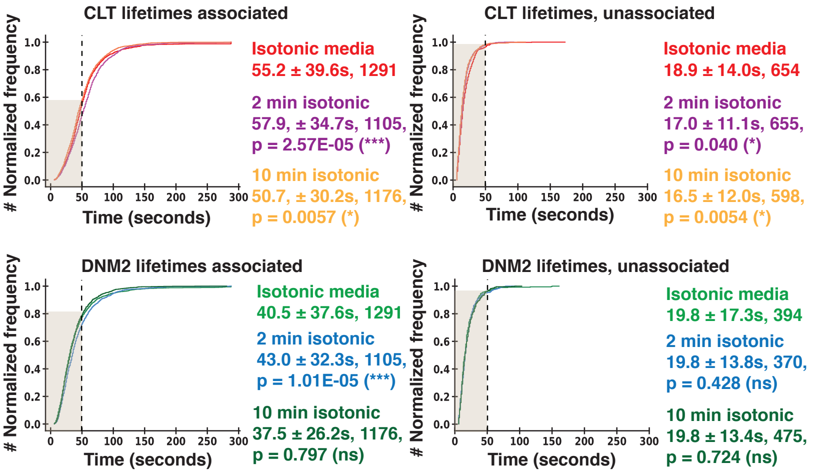

**B**

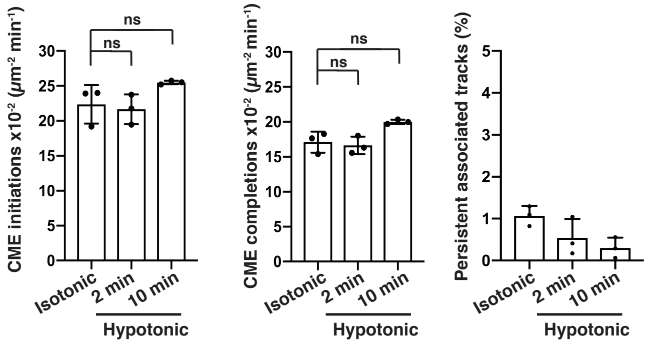

| | Mean initiation rate ± SD (100 $\mu m^2$*min$^{-1}$) | Mean completion rate ± SD (100 $\mu m^2$*min$^{-1}$) | Mean persistent tracks (%) ± SD |
|---|---|---|---|
| Isotonic | 19.19 ± 2.75 | 17.1 ± 1.51 | 1.1 ± 0.23 |
| Isotonic, 2 min | 19.46 ± 2.14 | 16.64 ± 1.26 | 0.54 ± 0.45 |
| Isotonic, 10 min | 25.22 ± 0.25 | 20 ± 0.33 | 0.30 ± 0.24 |

**C** Hypotonic media, 225 mOsm

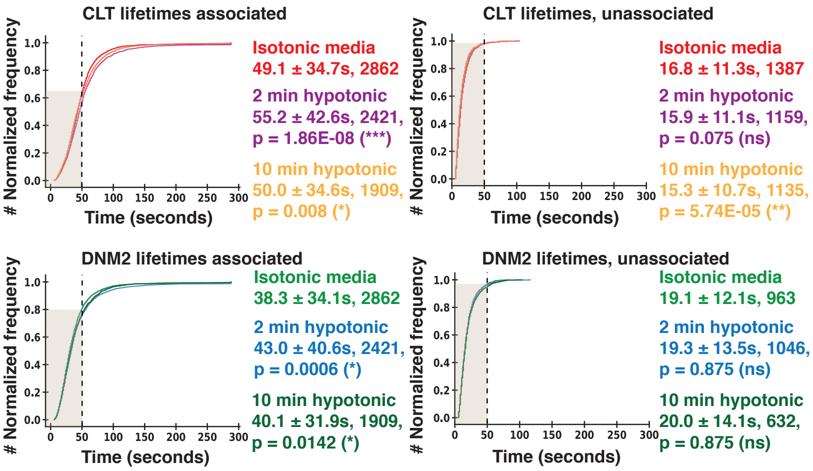

**D**

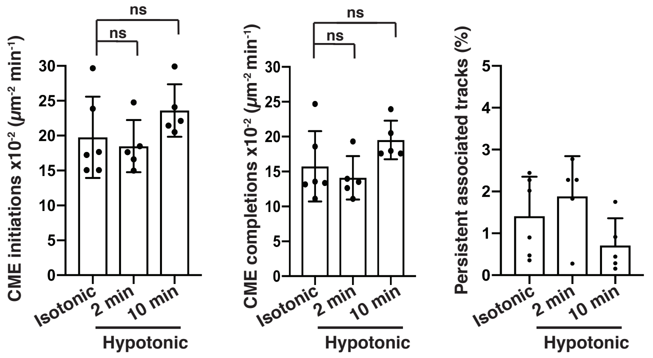

| | Mean initiation rate ± SD (100 $\mu m^2$*min$^{-1}$) | Mean completion rate ± SD (100 $\mu m^2$*min$^{-1}$) | Mean persistent tracks (%) ± SD |
|---|---|---|---|
| Isotonic | 19.76 ± 5.82 | 15.75 ± 5.03 | 1.41 ± 0.94 |
| 225 mOsm, 2 min | 18.50 ± 3.73 | 14.10 ± 3.12 | 1.89 ± 0.96 |
| 225 mOsm, 10 min | 23.61 ± 3.76 | 19.52 ± 2.76 | 0.71 ± 0.65 |

**E** Hypotonic media, 150 mOsm

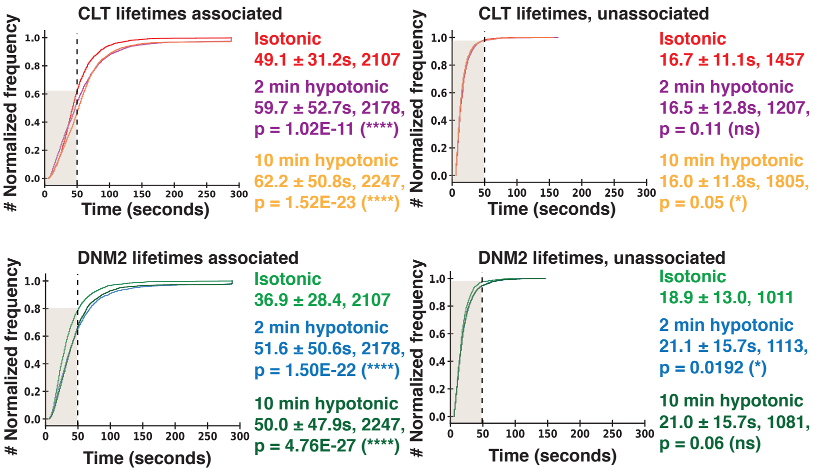

**F**

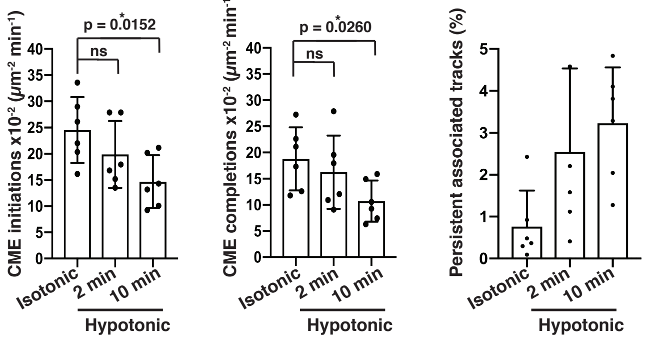

| | Mean initiation rate ± SD x10$^{-2}$ ($\mu m^2$*min$^{-1}$) | Mean completion rate ± SD x10$^{-2}$ ($\mu m^2$*min$^{-1}$) | Mean persistent tracks (%) ± SD |
|---|---|---|---|
| Isotonic | 24.53 ± 6.29 | 18.78 ± 6.03 | 0.77 ± 0.86 |
| 150 mOsm, 2 min | 19.87 ± 6.39 | 16.24 ± 7.02 | 2.55 ± 1.99 |
| 150 mOsm, 10 min | 14.70 ± 5.00 | 10.72 ± 3.92 | 3.23 ± 1.34 |

**G** Hypotonic media, 75 mOsm

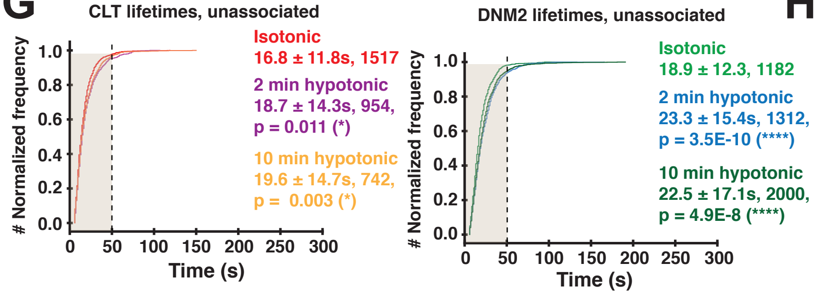

**H**

| | Mean initiation rate ± SD x10$^{-2}$ ($\mu m^2$*min$^{-1}$) | Mean completion rate ± SD x10$^{-2}$ ($\mu m^2$*min$^{-1}$) | Mean persistent tracks (%) ± SD |
|---|---|---|---|
| Isotonic | 22.18 ± 3.46 | 17.75 ± 3.43 | 0.65 ± 0.30 |
| 75 mOsm, 2 min | 10.28 ± 1.48 | 5.32 ± 1.72 | 19.17 ± 9.21 |
| 75 mOsm, 10 min | 9.54 ± 2.05 | 5.82 ± 2.07 | 11.45 ± 9.66 |

### Figure S8

A

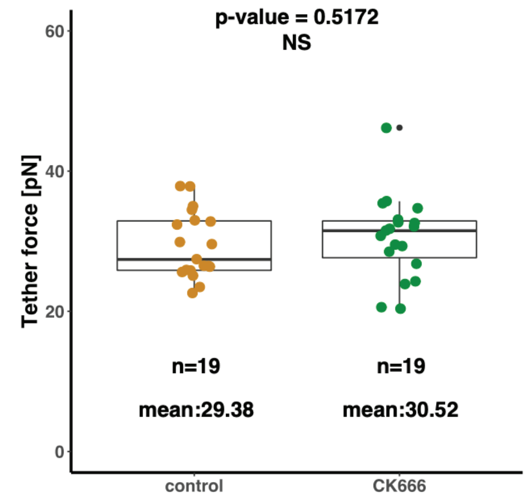

B

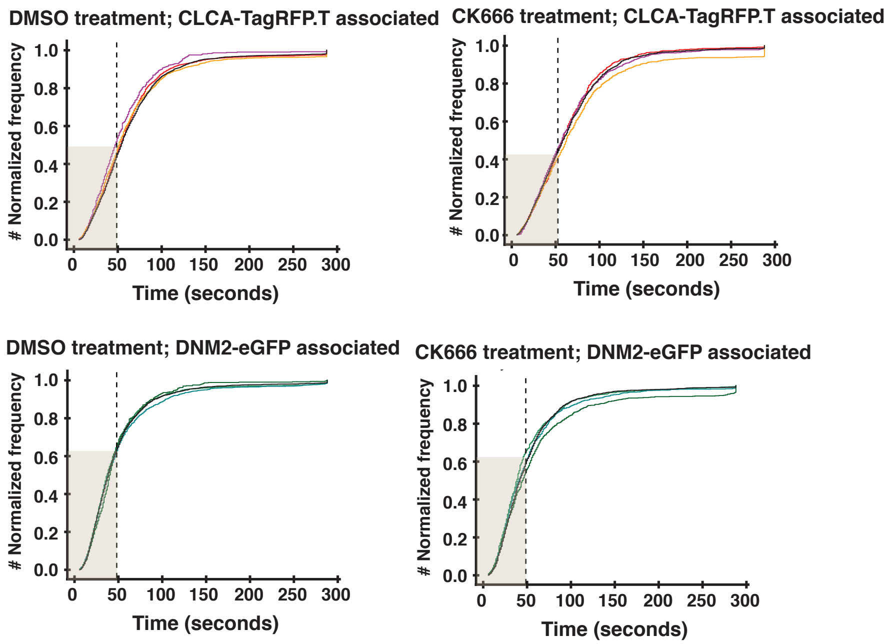

C

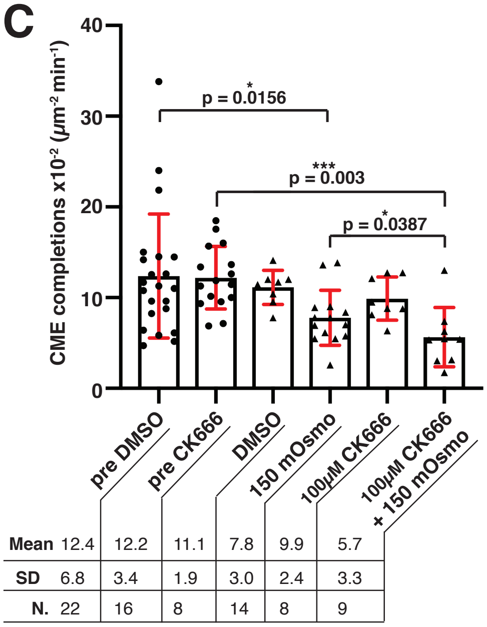

D

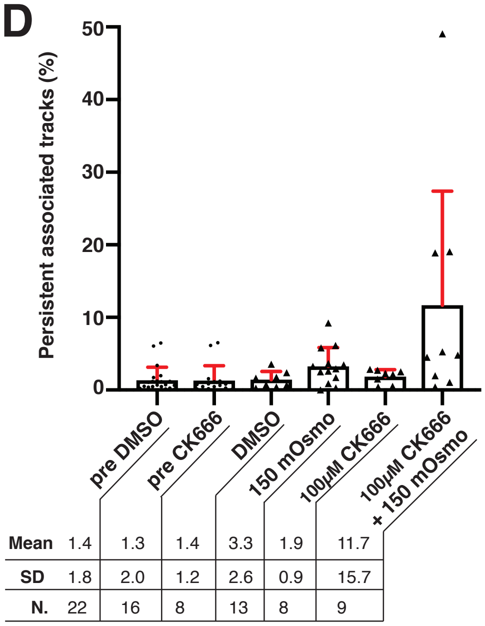

F

| | Mean height (nm) | SD (nm) | N. of values | Mean width (nm) | SD (nm) | N. of values |
|---|---|---|---|---|---|---|
| DMSO | 98 | 21 | 154 | 177 | 30 | 154 |
| 100µM CK666 | 106 | 27 | 158 | 177 | 28 | 158 |
| 100µM CK666 + 150mOsmo | 96 | 24 | 159 | 171 | 32 | 159 |

E

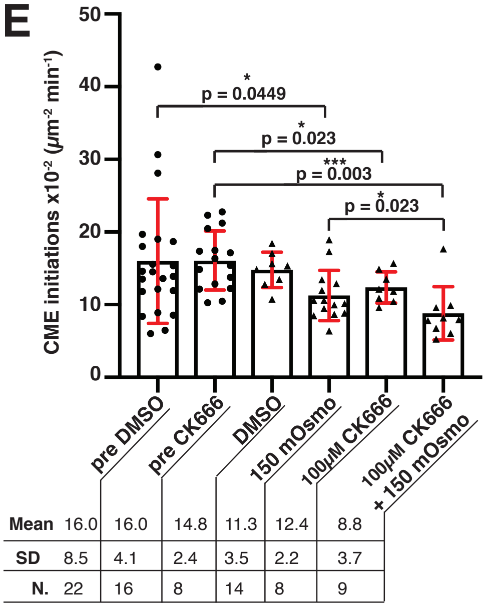

G

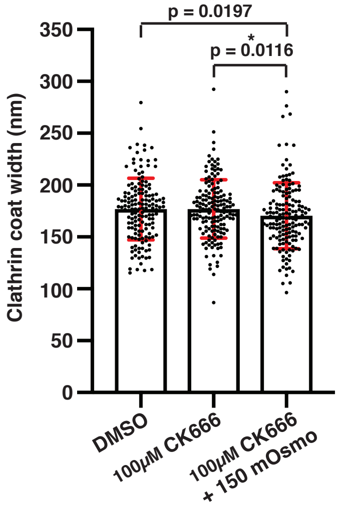

H

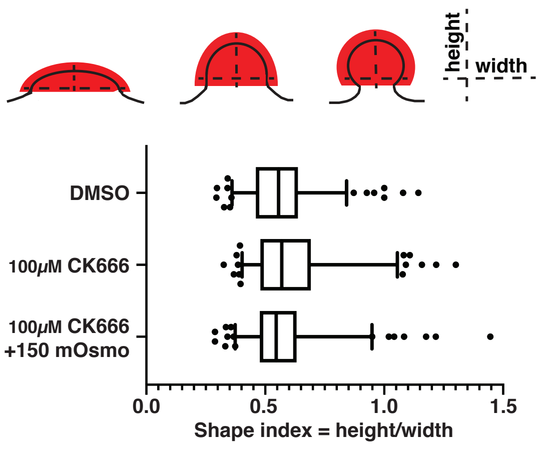

### Figure S9

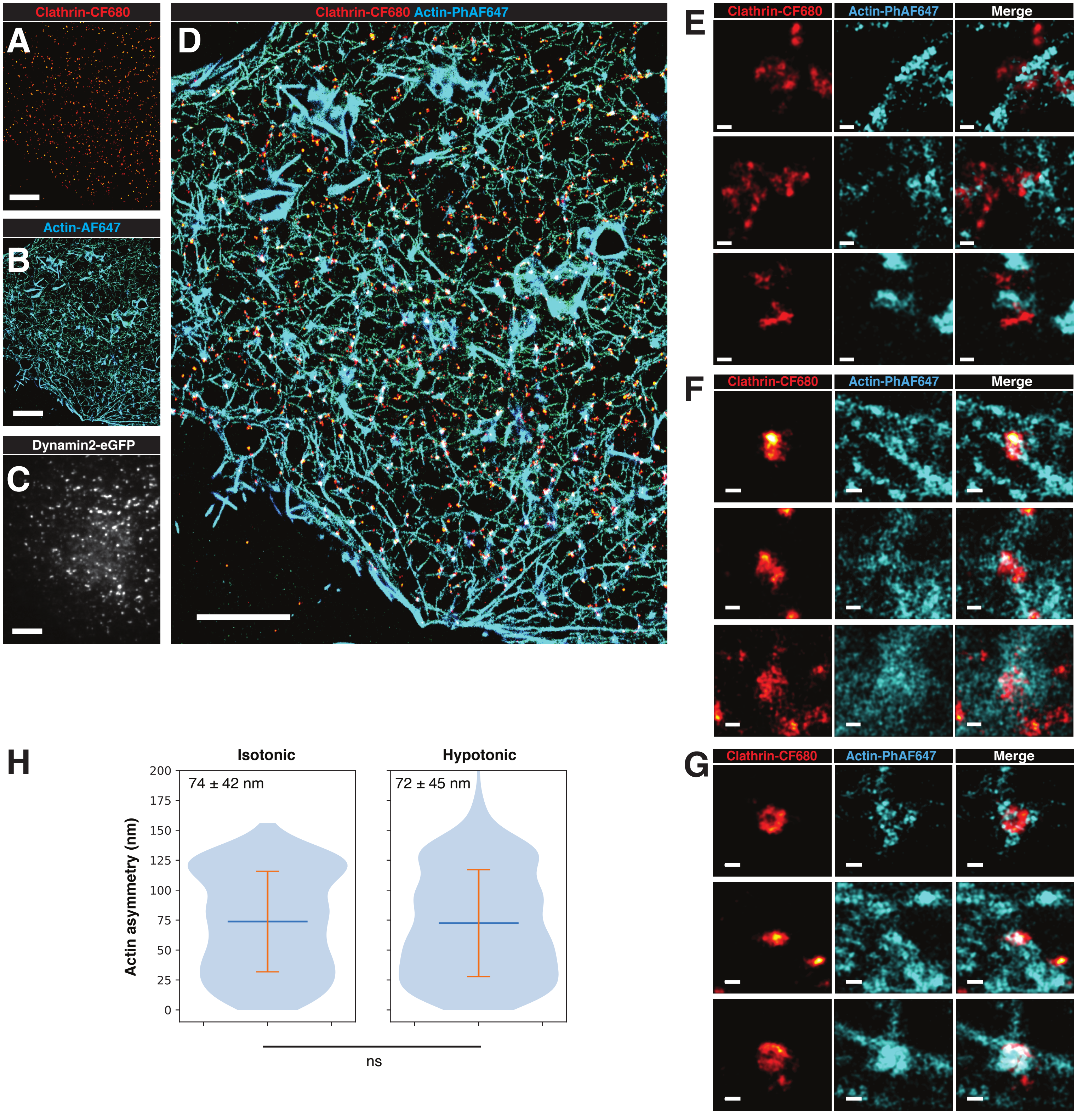

Clathrin-CF680
A
Actin-AF647
B
Dynamin2-eGFP
C
Clathrin-CF680 Actin-PhAF647
D
E
Clathrin-CF680
Actin-PhAF647
Merge
F
Clathrin-CF680
Actin-PhAF647
Merge
G
Clathrin-CF680
Actin-PhAF647
Merge
H
Isotonic
Hypotonic
74 ± 42 nm
72 ± 45 nm
Actin asymmetry (nm)
200
175
150
125
100
75
50
25
0
ns

### Figure S10

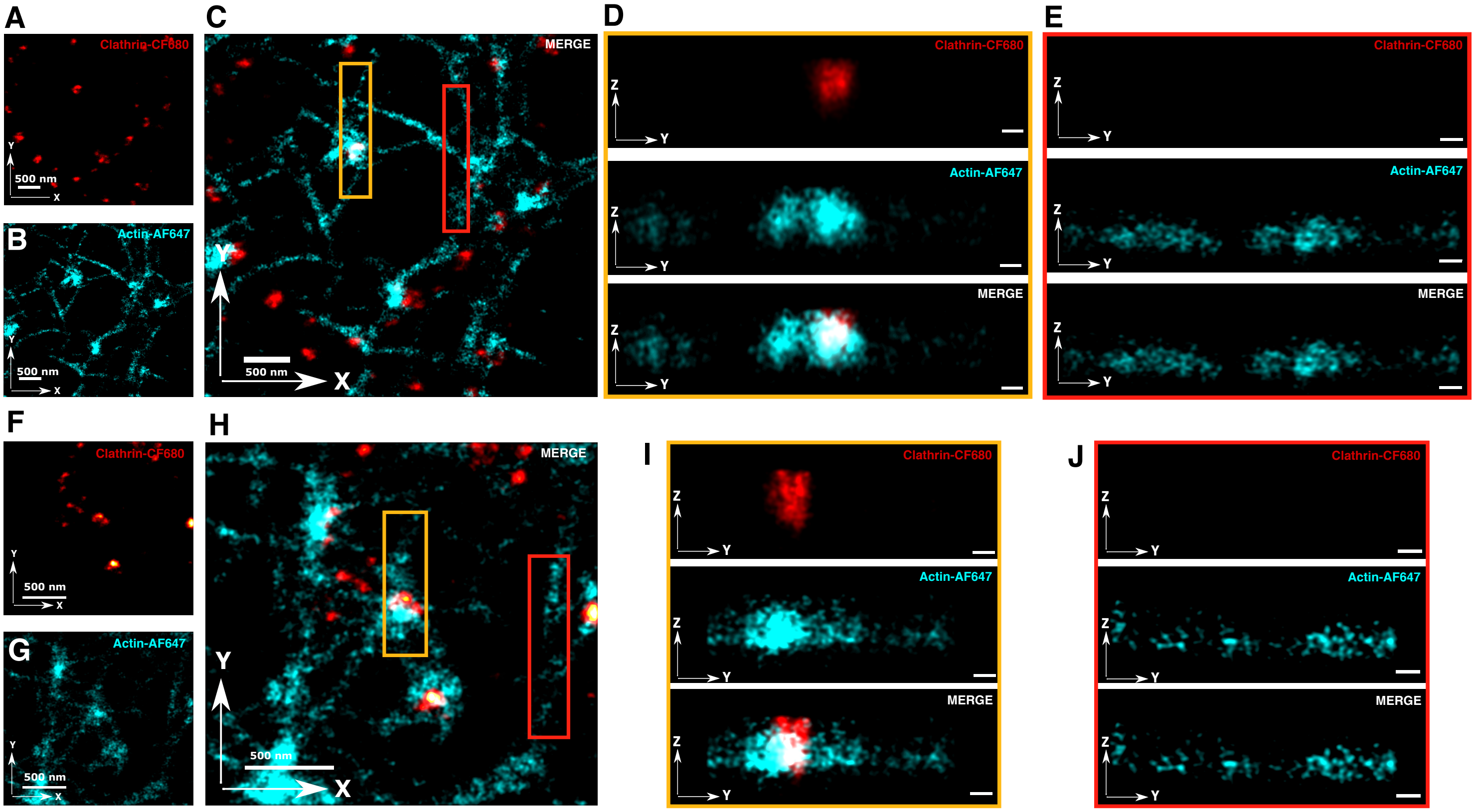
A
Clathrin-CF680
500 nm
B
Actin-AF647
500 nm
C
MERGE
500 nm
D
Clathrin-CF680
Actin-AF647
MERGE
E
Clathrin-CF680
Actin-AF647
MERGE
F
Clathrin-CF680
500 nm
G
Actin-AF647
500 nm
H
MERGE
500 nm
I
Clathrin-CF680
Actin-AF647
MERGE
J
Clathrin-CF680
Actin-AF647
MERGE

**Figure S1: Three-dimensional resolution of STORM set up.** Overlayed localization distributions of 24 representative single molecules from 3 different samples. Gaussian fits (red curves) gave standard deviations of 10 nm in-plane for the *XY* directions (left and middle plot) and 19 nm in-depth for the *Z* direction (right plot).

**Figure S2: Representative examples x-y and x-z STORM image projections to visualize the different appearance of actin at CME sites compared to cortical actin. (A)** and **(F)** X-y projections of super-resolved clathrin coats (red). Regions of interest (ROI) selected from STORM image in Figure 1A. **(B)** and **(G)** Corresponding super-resolved x-y projections of actin (cyan) to ROIs in (A) and (F). (**C)** and **(H)** Dual-color x-y projection from STROM images of clathrin (A, F, red) and actin (B, G, cyan). The yellow square inset shows a clathrin coated pit (CCP) with associated actin. The red squared inset shows actin at the cortex without a clathrin coated structure (CCS). **(D) - (J)** X-z STORM projections of red and yellow squared inset from C and H. The upper image shows clathrin (D, I, red) or no clathrin (E, J), the middle image shows actin (rainbow) and the lower image the merge of both images. In example images D and I actin is enriched and builds up at the CCP, therefore is distinguishable from the actin that extends away from the CCP. In examples E and J actin is not associated with a CCS and does not show a specific enrichment and accumulation as compared to the image in D and I. **(A) - (J)** All scale bars are 100 nm.

**Figure S3: Representative examples of x-y STORM image projections of single and double clathrin structures. (A)** Electron micrograph (image 1) shows a single clathrin coated pit in a SK-MEL-2 cell acquired by platinum replica EM. Image 2 shows a clathrin coated pit (red) resolved by STORM of the same size as the pit in the electron micrograph. X-y STROM projections of diverse single clathrin coated structures and pits (image 2 to image 21) in SK-MEL-2 cells. **(B)** Electron micrograph (image 1) shows a double clathrin coated pit structure in a SK-MEL-2 cell acquired by platinum replica EM. Image 2 shows a double clathrin coated pit structure (red) resolved by STORM of the same size as the double pit in the electron micrograph. X-y STROM projections of diverse double clathrin coated structures and pits (image 2 to image 21) in SK-MEL-2 cells. The two clathrin coated structures that are forming a double structure are highlighted with a white dashed circle. **(A) - (B)** Scale bars all 100 nm.

**Figure S4: Representative examples of ambiguous clathrin data, overview of 3D measurement methods for super-resolved clathrin and actin, and representative x-y projections.**

**(A)** Representative examples of immunolabeled clathrin structures that cannot be identified as individual clathrin coated events. STORM images show x-y projection of clathrin structures. First two columns show structures that do not show a clear round or elliptical shape. Third image column shows clathrin structures that extend over the entire region of interest. Fourth image column shows very small punctate structures. **(B)** Illustration of method to obtain actin and clathrin height and extent of coverage from super-resolved CCP images. x-z projection of a merged STORM image of clathrin and actin with its corresponding normalized pixel intensity histogram projected onto the z-axis. The black line marks the $30^{th}$ percentile of the z height histogram, which we used to calculate the height of clathrin and actin. We found that this metric, the "full z width at $30^{th}$ percentile max," was more reliable than the full width at half max. These raw intensity profiles are noisy based on the image quality, which depends on the fluorescent labeling quality of the target structure. Without filtering the profile, the measured height of clathrin and actin are underestimated. Numbers correspond to positions used to calculate height (in z) of clathrin (1. and 2.) and actin (4. and 5.) and their average positions (3. and 6.). Lower panel shows the same STORM image after a mean filter was applied to reduce noise in the histogram of z-position to improve the reliability of the clathrin and actin height readout. We measured the coverage of clathrin and actin profiles by subtracting the upper position of actin from the lower position of clathrin. **(C)** x-z projected STORM image of a clathrin coat with its corresponding normalized pixel intensity histogram projected onto the x-axis unfiltered. Due to the image noise the measurement of clathrin coat width misses most of the image. The right image shows the improved width measurement of the same clathrin coat after application of a median filter to smooth the histogram. All scale bars are 100 nm. **(D)** x-y STORM image projections corresponding to the x-z projections in Fig. 1E. **(E)** Example of asymmetrically organized actin around a clathrin-coated pit. Normalized pixel intensity histograms of clathrin (red) and actin (blue) from x-z and y-z projections. Measurement is the distance of the peak actin signal from the center of the clathrin coat. Actin is asymmetrically organized since the distance from the x-axis to the center is large compared to the distance to the y-axis. **(F)** Violin plot of actin asymmetry values in isotonic condition for three endocytic stages. Differences are not statistically significant based on Mann-Whitney test. N = 289 (shallow), 405 (U-shape), and 602 (omega shape).

**Figure S5: Clathrin and actin scatter plots of geometric parameters obtained from xz and yz STORM image projections. (A) - (D)** Data set of clathrin and actin parameters when cells were treated with isotonic media, n = 731. **(A)** Scatter plot of shape indices (blue dots) and regression line (red) with an $r^2 = 0.92$. **(B)** Scatter plot of clathrin height (red) and regression line (black) with an $r^2 = 0.99$. **(C)** Scatter plot of clathrin width (red) and regression line (black) with an $r^2$ =

0.93. **(D)** Scatter plot of actin height (green) and regression line (red) with an $r^2 = 0.99$. **(E) - (H)** Data set of clathrin and actin parameters when cells were treated with hypotonic media. n = 473 **(E)** Scatter plot of shape indices (blue dots) and regression line (red) with an $r^2 = 0.94$. **(F)** Scatter plot of clathrin height (red) and regression line (black) with an $r^2 = 0.99$. **(G)** Scatter plot of clathrin width (red) and regression line (black) with an $r^2 = 0.95$. **(H)** Scatter plot of actin height (green) and regression line (red) with an $r^2 = 0.99$.

**Figure S6: STORM images of N-WASP at clathrin-coated pits in SK-MEL-2 cells.** Schematic of clathrin coat side profiles shows N-WASP localization. Merged STORM images show spatial localization of N-WASP-AF647 (rainbow) at clathrin coats (red) in x-z projections. **(A)** N-WASP localized at the base of clathrin coats. **(B)** N-WASP localized all over clathrin coats. Scale bars are 100 nm.

**Figure S7: Hypotonic media treatment of SK-MEL-2 cells endogenously expressing CLTA-TagRFP-T$^{EN}$ and DNM2-eGFP$^{EN}$. (A), (C)** and **(E)** Normalized cumulative distribution data for CLTA-TagRFP-T$^{EN}$ and DNM2-eGFP$^{EN}$ fluorescence lifetimes when associated with or not associated with each other. Lifetimes were recorded under isotonic media conditions and then recorded after 2 min or 10 min of media exchange. Plot legend provides, respectively: mean lifetime ± SD, number of tracks, p-value determined in Kolmogorov-Smirnov test with significance compared to the control. **(B), (D)** and **(F)** show CME initiation rate, completion rate and percentage of persistent tracks plotted for sites in which DNM2-eGFP$^{EN}$ and CLTA-TagRFP-T$^{EN}$ were associated with each other. Tables below show the corresponding statistics. Mann-Whitney statistical test was used to compare CME initiation rates and completion rates. **(A)** and

**(B)** Control conditions in which only isotonic media exchange was performed. n = 3-4 (cells) for each condition. Experiments were repeated 3 times. **(C)** and **(D)** Media were exchanged from isotonic to 225 mOsm hypotonic. For each of the following conditions: control, 2 min, or 10 min after media exchange, n = 6 (cells). **(E)** and **(F)** Media were exchanged from isotonic to 150 mOsm hypotonic. For each condition of the following conditions: control, 2 min and 10 min after media exchange, n = 6 (cells). **(G)** Unassociated tracks in 75 mOsm hypotonic media. **(C) – (G)** Experiments were repeated 5 – 6 times. **(H)** Statistics for CME initiation rates, completion rates and percentage of persistent tracks for CLTA-TagRFP-$T^{EN}$ associated with DNM2-$eGFP^{EN}$ corresponding to Figure 2 F – H.

**Figure S8: Effects of hypotonic media and concomitant CK666 treatment on membrane tether force and CME progression. (A)** Membrane tether force measurements from atomic force microscopy in cells treated with DMSO (left) or 100 µM CK666 (right). n=19 cells in each condition. 4 biological replicates; minimum 3 tethers per cell. Tether force was measured 2-19 mins after addition of CK666. p>0.5 by t-test. Boxplot shows median with interquartile range. **(B)** Normalized fluorescence lifetime cumulative distribution data for CLTA-TagRFP-$T^{EN}$ and associated DNM2-$eGFP^{EN}$ for CK666 concentrations of 25 µM, 50 µM and 100 µM. Respective controls were 0.5% v/v, 1% v/v and 2% v/v DMSO. Fluorescence lifetimes were acquired after 2 minutes of treatment. Experiments were repeated 2-3 times. Kolmogorov-Smirnov statistical test was used. Statistics are provided in Table S2. **(C)** Corresponding CME completion rates for CLTA-TagRFP-$T^{EN}$ associated with DNM2-$eGFP^{EN}$ in Figure 3 B - F. Barplots show mean ± SD **(D)** Percentage of persistent tracks for CLTA-TagRFP-$T^{EN}$ associated with DNM2-$eGFP^{EN}$ for corresponding imaging data in Figure 3 B - F. Barplots show mean ± SD **(E)** Corresponding CME initiation rates for CLTA-TagRFP-T$^{EN}$ associated with DNM2-eGFP$^{EN}$ in Figure 3 B - F. Barplots show mean ± SD **(F)** Statistics for clathrin coat height and width for data in Figure 3 H when cells were treated with 2% v/v DMSO, 100 μM CK666 and 100 μM CK666 in 150 mOsm hypotonic media. **(G)** Plotted clathrin coat width in nm for data in Figure 3 H. Barplots show mean ± SD. **(H)** Ratio of clathrin coat height to width for data in Figure 3 H and Figure S3 G. **(C), (E)** and **(G)** Mann-Whitney statistical test was used.

**Figure S9: Effect of hypotonic media treatment on actin organization. (A)** STORM image of CF-680 immunolabeled clathrin-coated structures (red) on the ventral cell surface when 75 mOsm hypotonic media was applied for 5 min. **(B)** Same area of the cell as in (A) showing the STORM image of phalloidin-AF647 labeled actin cytoskeleton (rainbow). **(C)** Same area of the cell as in (A) showing the corresponding DNM2-eGFP$^{EN}$ conventional microscopy image. **(D)** Merged clathrin and actin STORM images from (A) and (B). Color bar shows actin position in the z-dimension. **(A) – (D)** All scale bars are 5 μm. **(E) - (G)** STORM x-y projection images corresponding to images in Fig. 4 B. (E) corresponds to the shallow invaginations. (F) corresponds to the U-shaped invaginations. (G) corresponds to the omega-shaped invaginations. All scale bars are 100 nm. **(H)** Violin plot of actin asymmetry values in isotonic and 75 mOsm hypotonic conditions. Differences are not statistically significant based on Mann-Whitney test. Data for isotonic conditions are also reported in Fig S1. Hypotonic condition: N = 1090.

**Figure S10: Representative examples x-y and y-z STORM image projections to visualize the different appearance of actin at CME sites compared to cortical actin in cells treated with**

**hypotonic media. (A)** and **(F)** Example regions of interest showing X-y projections of super-resolved clathrin coats (red). Regions of interest (ROI) selected from STORM image in Supp Figure 5C. **(B)** and **(G)** Corresponding super-resolved x-y projections of actin (cyan) to ROIs in (A) and (F). (**C)** and **(H)** Dual-color x-y projection from STROM images of clathrin (A, F, red) and actin (B, G, cyan). The yellow square inset shows a clathrin coated pit (CCP) with associated actin and extends along the cell cortex. The red squared inset shows actin at the cortex without a clathrin coated structure (CCS). **(D) - (J)** Y-z STORM projections of red and yellow squared insets from C and H. The upper image shows clathrin (D, I, red) or no clathrin (E, J), the middle image shows actin (cyan) and the lower image the merge of both images. In example images D and I actin is enriched and builds up at the CCP, therefore is distinguishable from the actin that extends away from the CCP. In examples E and J actin is not associated with a CCS and does not show a specific enrichment and accumulation as compared to the image in D and I. **(A) - (C)** and **(F) - (H)** All scale bars are 500 nm. **(D), (E)**, **(I) - (J)** All scale bars are 100 nm.

**Table S1: Lifetimes of endocytic tracks in isotonic and hypotonic conditions.** P values determined by Kolmogorov-Smirnov test.

**Table S2: Lifetimes of endocytic tracks in cells treated with the small molecule inhibitor CK666.** P values determined by Kolmogorov-Smirnov test.

**Table S3: Lifetimes of endocytic tracks in cells in hypotonic conditions and treated with the small molecule inhibitor CK666.** P values determined by Kolmogorov-Smirnov test.

**Supplementary movie 1:** Movie of CLTA-TagRFP-$T^{EN}$ (magenta) and DNM2-$eGFP^{EN}$ (green) in SK-MEL-2 cells in isotonic media.

**Supplementary movie 2:** Movie of CLTA-TagRFP-$T^{EN}$ (magenta) and DNM2-$eGFP^{EN}$ (green) in SK-MEL-2 cells after 2 minutes of media exchange to 75 mOsm media.

**Supplementary movie 3:** Movie of CLTA-TagRFP-$T^{EN}$ (magenta) and DNM2-$eGFP^{EN}$ (green) in SK-MEL-2 cells after 10 minutes of media exchange to 75 mOsm media.

**Supplementary movie 4:** Movie of CLTA-TagRFP-$T^{EN}$ (magenta) and DNM2-$eGFP^{EN}$ (green) in SK-MEL-2 cells in isotonic media before treatment with CK666 and 150 mOsm hypotonic media.

**Supplementary movie 5:** Movie of CLTA-TagRFP-$T^{EN}$ (magenta) and DNM2-$eGFP^{EN}$ (green) in SK-MEL-2 cells in 150 mOsm hypotonic media with 100 μM CK666. Movie acquisition started 2 minutes after media exchange.

**Data sheet 1:** Data set of clathrin and actin values of height, width, shape index (aspect ratio) and maximum position in the Xz - and Yz - projections. Control data set. Cells were under isotonic media conditions.

**Data sheet 2:** Clathrin height and width under isotonic media conditions, CK 666 and 150 mOsm hypotonic media plus CK 666 treatment.

**Data sheet 3:** Data set of clathrin and actin values of height, width, shape index (aspect ratio) and maximum position in the Xz - and Yz - projections. Cells were under 75 mOsm hypotonic media conditions.
